# Enchained growth and cluster dislocation: a possible mechanism for microbiota homeostasis

**DOI:** 10.1101/298059

**Authors:** Florence Bansept, Kathrin Schumann-Moor, Médéric Diard, Wolf-Dietrich Hardt, Emma Slack, Claude Loverdo

## Abstract

Immunoglobulin A is a class of antibodies produced by the adaptive immune system and secreted into the gut lumen to fight pathogenic bacteria. We recently demonstrated that the main physical effect of these antibodies is to enchain daughter bacteria, i.e. to cross-link bacteria into clusters as they divide, preventing them from interacting with epithelial cells, thus protecting the host. These links between bacteria may break over time. We study several models using analytical and numerical calculations. We obtain the resulting distribution of chain sizes, that we compare with experimental data. We study the rate of increase in the number of free bacteria as a function of the replication rate of bacteria. Our models show robustly that at higher replication rates, bacteria replicate before the link between daughter bacteria breaks, leading to growing cluster sizes. On the contrary at low growth rates two daughter bacteria have a high probability to break apart. Thus the gut could produce IgA against all the bacteria it has encountered, but the most affected bacteria would be the fast replicating ones, that are more likely to destabilize the microbiota. Linking the effect of the immune effectors (here the clustering) with a property directly relevant to the potential bacterial pathogeneicity (here the replication rate) could avoid to make complex decisions about which bacteria to produce effectors against.

**Author Summary:** Inside the organism, the immune system can fight generically against any bacteria. However, the lumen of the gut is home to a very important microbiota, so the host has to find alternative ways to fight dangerous bacteria while sparing beneficial ones. While many studies have focused on the complex molecular and cellular pathways that trigger an immune response, little is known about how the produced antibodies act once secreted into the intestinal lumen. We recently demonstrated that the main physical effect of these antibodies is to cross-link bacteria into clusters as they divide, preventing them from interacting with epithelial cells, thus protecting the host. These links between bacteria may break over time. Using analytical and numerical calculations, and comparing with experimental data, we studied the dynamics of these clusters. At higher replication rates, bacteria replicate before the link between daughter bacteria breaks, leading to growing cluster sizes, and conversely. Thus the gut could produce IgA against all the bacteria it has encountered, but the most affected bacteria would be the fast replicating ones, that are more likely destabilize the microbiota. Studying the mechanisms of the immune response may uncover more such processes that enable to target properties hard to escape through evolution.

## Introduction

The digestive system has a large surface area[1][2] covered by a single layer of epithelial cells, essential for nutrient absorption, but also a gateway for many pathogens. Contrary to the inside of the body, where the presence of any bacteria is abnormal, the lumen of the digestive system is home to a very important microbiota. These microbiota bacteria are present in extremely high densities[3]. Bacteria are necessary to break down and absorb certain nutrients, and can compete against potentially pathogenic intruders[4]. Inside the organism, the immune system can fight generically against any bacteria. However, in the digestive system, the host has to find alternative ways to fight dangerous bacteria while sparing beneficial ones. As closely related bacteria (e.g. *Salmonella* spp. and commensal *E. coli)* can show highly variable behaviors in the intestine, identifying which bacteria are good or bad is challenging. Besides, the over-growth of any type of bacteria, even those that do not cause acute pathology, can impair the functionality of the microbiota. Thus the host needs mechanisms to maintain the gut microbiota homeostasis.

The adaptive response is the only strong handle that the host has on directly controlling microbiota composition at the species level[5][6]. The main effector of the adaptive immune response in the digestive system is secretory IgA, an antibody. sIgA specifically bind to targets that the organism has already encountered and can be elicited by vaccination. It was observed more than 40 years ago that this prevents infection by pathogenic bacteria such as salmonella[7]. Many studies have focused on the complex molecular and cellular pathways that trigger an immune response on the host side of the digestive surface[8]. However, we are only just beginning to understand by which physical mechanisms the immune effectors act once secreted into the intestinal lumen, which are crucial for the control of both commensal and pathogenic bacteria.

The influence on bacteria dynamics of abiotic factors such as the flow in the gut has recently started being quantitatively studied [9, 10].

We have shown that mice vaccinated with inactivated *Salmonella* Typhimurium do produce specific sIgA which bind to *S*.Typhimurium, but this neither kills them nor prevents them from reproducing[11][12]. The initial colonization of the intestinal lumen by *S*.Typhimurium is in fact unchanged in either kinetics or magnitude in vaccinated animals. These mice are nevertheless protected against pathogen spread from the gut lumen to systemic sites like lymph nodes, liver or spleen. A classic idea in immunology is that, as one antibody has several binding sites, antibodies aggregate bacteria when they collide into each other. But this effect would be negligible at realistic densities of a given bacterium in the digestive system, simply due to very long typical encounter times between bacteria recognized by the same sIgA (see section, in appendix). We have shown that actually, the main effect is that upon replication, daughter bacteria remain attached to one another by sIgA, driving the formation of clusters derived from a single infecting bacterium[12]. This 11enchained growth1, is effective at any bacterial density. Clustering has physical consequences: the produced clusters do not come physically close to the epithelial cells. And as interaction with the epithelial cells is essential for *S*.Typhimurium virulence, this is sufficient to explain the observed protective effect.

If sIgA was perfectly sticky, we would expect all bacteria to be in clusters of ever increasing size. In these experiments, despite observing *S*.Typhimurium clusters in the presence of sIgA, there are still free *S*.Typhimurium, and small clusters. One possibility would be that not all bacteria are coated with sIgA. But in these experiments, it has been demonstrated that they are (extended figure 2c of[12]). Indeed, a gram of digestive content contains at most 10^1,^ bacteria, and typically 50 micrograms or more of sIgA[13], of molecular mass of about 385kD. This leads to about 800 sIgA per bacteria. sIgA may not be all bound to bacteria, and sIgA for different specific antigens may be produced in proportions not matching the proportions of antigens present in the digestive system, so that not all bacteria are coated with 800 sIgA. Nevertheless, most bacteria already encountered by the organism will be coated with many sIgA, and thus the cluster size is not limited by the number of available sIgA. Another possibility is that the sIgA-mediated links break. Such breaking has been demonstrated to be dependent on the applied forces in related systems[14][15]. As there is shear in the digestive system, because mixing is needed for efficient nutrients absorption, it is plausible that links break over time.

Small clusters are linear chains of bacteria, bound by sIgA, with these links being broken over time by the forces induced by the flow. As bacteria are similar to each other, it is, at another scale, analogous to other physical systems[16] such as polymers breaking under flow[17]. The main difference is that these chains grow by bacterial replication. Growth and fragmentation are competing effects, and the modelling of these chains can be viewed as statistical physics, to predict their length distribution, whether there is a typical chain length, or if large chains of ever-increasing length dominate the distribution, and how the growth in number of free bacteria depends on the bacterial replication rate.

This could have very important biological consequences. To illustrate this point, let us consider a simplified model: bacteria remain enchained by sIgA when they grow (replication time *τ*_*div*_), and this link between 2 bacteria breaks at a specific time *τ*_*break*_ (although this latter hypothesis is not realistic, we make it for now for the sake of simplicity). If *τ*_*div*_ > *τ*_*break*_, then when a bacterium divides, it forms a 2-bacteria cluster, which dislocates into 2 free bacteria before the next replication steps, so that the bacteria remain in the state of free or 2-bacteria clusters and there are no larger clusters. If *τ*_*div*_ < *τ*_*break*_, when a bacterium divides, it forms a 2-bacteria cluster, which becomes a 4 bacteria cluster before the first link breaks, so there cannot be free bacteria. In this model, the fast-growing bacteria are selectively targeted by the action of the immune system. The immune system does not need to sense which bacteria are growing faster, it only has to produce sIgA targeted to all the bacteria it has encountered, and bacteria with *τ*_*div*_ > *τ*_*break*_ are unaffected, whereas bacteria with *τ*_*div*_ < *τ*_*break*_ are trapped in clusters. That could be a simple physical mechanism to target the action of the immune system to the fast-growing bacteria which are destabilizing the microbiota, and thus to preserve microbiota homeostasis.

In the following, we present different plausible models of bacteria clusters dynamics, and the methods to study them. Then we give, for each model, the resulting dynamics and chain length distribution, before putting these results in perspective with experimental data. Eventually, we discuss the results. As some biological details are unknown, studying different models enables to show which key results are robust; and differences confronted to experimental data give some indications about which are the most likely.

## Models and methods

### Ethics statement

All animal experiments were approved by the legal authorities (licenses 223/2010, 222/2013 and 193/2016; Kantonales Veterinäramt Zürich, Switzerland) and performed according to the legal and ethical requirements.

### Experimental methods

We perform a new analysis on microscopy images that were produced for [12]. We analyzed images of cecal content in vaccinated mice for the early data points (4 and 5 hours) of experiments starting from a low inoculum (10^5^), to minimize the clustering from random encounters. Further details on our analysis can be found in appendix 7, as well as a brief description of the experiments from which the images were produced.

### Models and general methods

We consider low bacterial densities, so encounters between unrelated bacteria are negligible. Thus, we consider each free bacteria and each cluster of bacteria independently of the others. *Salmonella* are rod shaped bacteria, which replicate by dividing in two daughter bacteria at the middle of the longitudinal axis. Thus if the daughter bacteria remain enchained, they are linked to each other by their poles. With further bacterial replications, the cluster will then be a linear chain. This is consistent with experimental observations, in which clusters are either linear chains, with bacteria attached to one or two neighbors by their poles, or larger clusters which seem to be formed as bundles of such linear clusters (pannel A figure 1). Our aim is to model the dynamics of these chains.

**Figure 1:**
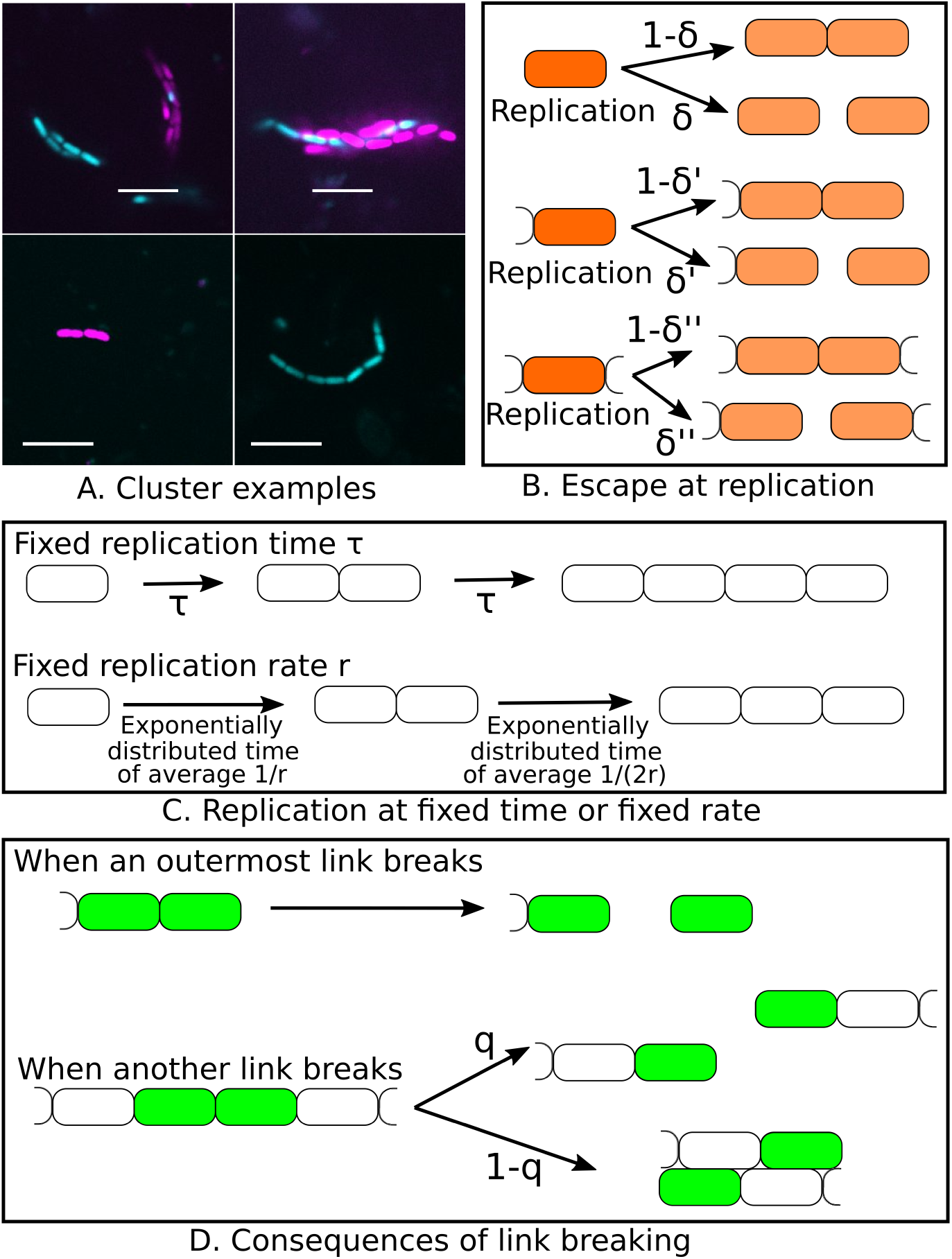
Bacterial cluster modeling. A. Representative experimental images of bacterial clusters in cecal content of vaccinated mouse at 5h post infection with isogenic GFP and mCherry expressing S.typhimurium (Experiments performed for [12]). The scale bar is 10*µm*. Top images: complex clusters made from bundles of linear clusters, which could be re-linked single chains (left) or formed from at least two independent clones (indicated by fluorescence, right). Bottom images: linear clusters which dynamics we aim to model. B. Potential bacterial escape at replication (in the base model, *δ* = *δ*′ = *δ*″). C. Fixed replication time or fixed replication rate (the latter is chosen for the base model). D. Consequences of link breaking. In the base model, *q* = 0.

A first element is the bacterial replication (see figure 1 C). One way to model it is to assume that bacteria replicate every *τ*_*div*_. Another way, that we will generally use, less realistic but easier for calculations, is to assume that there is a fixed replication rate *r*.

A second element is that when bacteria replicate, they may be able to escape enchainment (see figure 1 B), but likely with low probability (see discussion in section 2 in appendix). In general, we will take the limit with perfect enchainment upon replication (*δ* = *δ*′ = *δ*″ = 0).

A crucial element is the possibility for the links between bacteria to break. We usually assume that the breaking rate *α* is the same for all links and over time. We will also explore the case when the link breaking rate is force-dependent, in which case not all the links have the same breaking rate.

Another crucial element, is to model what happens when the chain breaks (see figure 1 D). If the subparts come in contact again at the same poles and get linked again, then this could simply be modeled by an effectively lower breaking rate. More likely, if the subparts come in contact again, they do so laterally, forming larger clusters of more complex shapes. Because in these clusters, most bacteria have more than two neighbors, and more contact surface, they are much less likely to escape. To simplify, we will consider that these clusters do not contribute anymore to releasing either free bacteria or linear chains. Thus when a link breaks, either the two subparts move sufficiently away and become two independent chains (probability *q*); or collide and become a more complex cluster which does not contribute anymore to both free bacteria and linear chains (probability 1*- q*). For simplicity, we consider that when an outermost link breaks, the single bacterium, more mobile, always escapes (*q*_*outermost*_ = 1), but that else *q* is length independent. The simplest values to study are either *q* = 0 or *q* = 1. As we will see, when we study the case in which *q* can take any value between 0 and 1, we find that the case *q* = 1 is qualitatively different from other values of *q*. Consequently, we will take *q* = 0 for the base model.

As digestive content leaves the digestive system, or the part of the digestive system under consideration, due to flow, we define *c* the loss rate of free bacteria, and *c*′ the loss rate of chains. We assume no deat. Bacterial death would break chains. It would thus have a similar effect to a larger breaking rate *α*. As free bacteria have more autonomous motility, enabling them to swim towards the epithelial cells, it is likely that *c*′≥*c*. We will usually take *c* = *c*′. Crucially, in this latter case, free bacteria, and all chains are lost at the same rate. The *c* value has a complex effect on stochastic quantities, such as the probability to have at least one chain of a given length. However, here we study the mean numbers of free bacteria and chains of different lengths, then the case with *c* = *c*′ is equivalent to *c* = *c*′ = 0, with all numbers of bacteria and chains multiplied by exp(-*ct*).

We start with the most basic model, with a replication rate *r*, bacteria perfectly bound upon replication (*δ* = *δ*′ = *δ*″ = 0), a fixed breaking rate per link *α*, and bacterial chains always binding into a more complex cluster when a link breaks (except for the outermost links) (*q* = 0). We then study variations of the model to test the robustness of the results: with an non-zero escape probability upon replication and *c* ≠*c*′; with a replication time *τ* instead of a replication rate *r*; with the possibility for chains to escape when an inner link breaks (*q* > 0); with a force-dependent breaking rate (see table below for a list of symbols).

We consider the beginning of the process, early enough so that the carrying capacity is far from reached, and thus the replication rate is constant. We do not consider generation of escape mutants which are not bound by IgA. We consider only the average numbers of free bacteria and linear chains of different lengths, and we do not count more complex clusters, as they do not contribute to free bacteria dynamics in our model. For each model, we write the equations for the derivative of these numbers with respect to time. With *N* the vector of the mean number of free bacteria, linear chains of length 2, 3, etc., these equations give the coefficients of the matrix *M*, such that *dN/dt* = *MN*. The results are obtained in part via analytical derivations and in part via numerical studies. The latter are obtained in Mathematica by numerically solving the eigensystem written for chains up to length *n*_*max*_, chosen large enough not to impact the results. In the long term limit, *N* (*t*) →*Ce*^*λt*^*P*, with *C* a constant, *λ* the largest eigenvalue of *M*, and *P* the corresponding eigenvector, normalized such that the sum of its components is equal to 1. *λ* is thus the long term growth rate of the free bacteria and the linear chains. For each model, we study how the growth of free bacteria - the ones which are capable of causing systemic infection[12] - which is *λ* in the steady state, depends on the bacterial replication rate. Besides, we obtain chain length distributions (the components *p*_*i*_ of *P)*, which could be compared to experimentally observed distributions.

## Recapitulation table of the symbols used

| **Base model** |  |
| --- | --- |
| $r$ | Bacterial replication rate |
| $\alpha$ | Breaking rate of the link between two bacteria |
| $n_i(t)$ | Number of linear chains of length $i$ at $t$ ($n_1$: free bacteria) |
| $\lambda$ | Largest eigenvalue of the matrix $M$,<br>which is the growth rate of the free bacteria in the steady state |
| $P$ | Eigenvector of components $p_i$ associated to $\lambda$,<br>normalized such as $\sum_{i=0}^{\infty} p_i = 1$. |
| **Model with bacterial escape and bacterial loss**<br>(all these parameters are taken as 0 in the base model) |  |
| $\delta$ | When a free bacterium replicates,<br>the probability that this will lead to 2 free bacteria |
| $\delta'$ | When a bacterium at the tip of a chain replicates,<br>the probability that the daughter bacterium at the exterior side escapes |
| $\delta''$ | When a bacterium replicates within a chain, the probability that the daughter bacterium will not be bond to each other, resulting to the chain breaking in two |
| $c$ | Loss rate for the free bacteria |
| $c'$ | Loss rate for the chains |
| **Model with fixed replication time** |  |
| $\tau$ | Time between one bacterial division and the next<br>(the bacterial growth rate is $r_{eff} = \log(2)/\tau$) |
| $\mathcal{N}$ | Largest eigenvalue of the matrix in this model. $\mathcal{N} = \exp(\lambda\tau)$ |
| **Model with linear chains independent after breaking** |  |
| $q$ | Probability that when a inner link of a chain breaks,<br>the two subparts become independent linear chains. In the base model, $q = 0$. |
| **Model with force-dependent breaking rate** |  |
| $\beta$ | A constant expressing the strength of the coupling between hydrodynamic forces<br>and link breaking. In the base model, $\beta = 0$. |

## Results

### Base model: replication rate, no bacteria escape upon replication, fixed breaking rate, *q* = 0

#### Equations

In the base model, bacteria have a replication rate *r*, daughters are perfectly bound upon replication, each link has a breaking rate *α*, and when a link which is not at a tip breaks, the resulting two chains of bacteria always bind into more complex clusters and thus do not contribute to free bacteria dynamics anymore (*q* = 0). With *n*_*i*_(*t*) the number of linear chains of length *i* as a function of time, (*n*_1_ is the number of free bacteria),

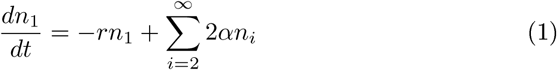

and for *i* ≥ 2,

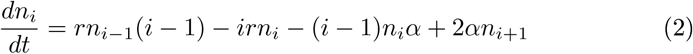

In a chain of length *i, i* bacteria may replicate, thus the total rate for one replication occurring in any bacterium of the chain is *ir*. This explains the terms *rn*_*i−1*_(*i* − 1) *− irn*_*i*_. Such a chain is made of *i* − 1 links, so the total breaking rate is (*i* − 1)*α*, explaining the term −(*i* − 1)*n*_*i*_*α*. When the link within a chain of length two breaks (rate *α* per chain), two free bacteria are released. For longer chains, there are two outermost links (on each side of the chain) which breaking releases one free bacterium and one one-bacterium shorter chain. This explains the term 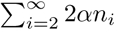 in equation (1) and the term 2*αn*_*i*+1_ in equation (2).

### Free bacteria growth rate as a function of the bacterial replication rate

Even for this simple version, the system of equations is hard to solve in the general case. We start by studying numerically the growth rate in the long term (the maximum eigenvalue *λ* of the matrix of coefficients *m*_*i,j*_ = *r*(*i* − 1)*δ*_*i−*1,*j*_*−irδ*_*i,j*_ − (*i −* 1)*αδ*_*i,j*_ + 2*α*(*δ*_*i*+1,*j*_ + *δ*_*i*,1_(1 *− δ*_*j*,1_ *− δ*_*j*,2_)), with *δ*_*i,j*_ the Kronecker symbol, which takes the value 1when *i* = *j*, and 0 otherwise), as a function of the replication rate (see figure 2A). The growth rate has a maximum for a finite replication rate, of the order of *α* (the link breaking rate): the higher the replication rate, the higher the potential for growth in the number of free bacteria, but when the replication rate becomes too large compared to the breaking rate, the bacteria get trapped in clusters, which break and re-attach in more complex clusters from which independent bacteria cannot escape.

**Figure 2:**
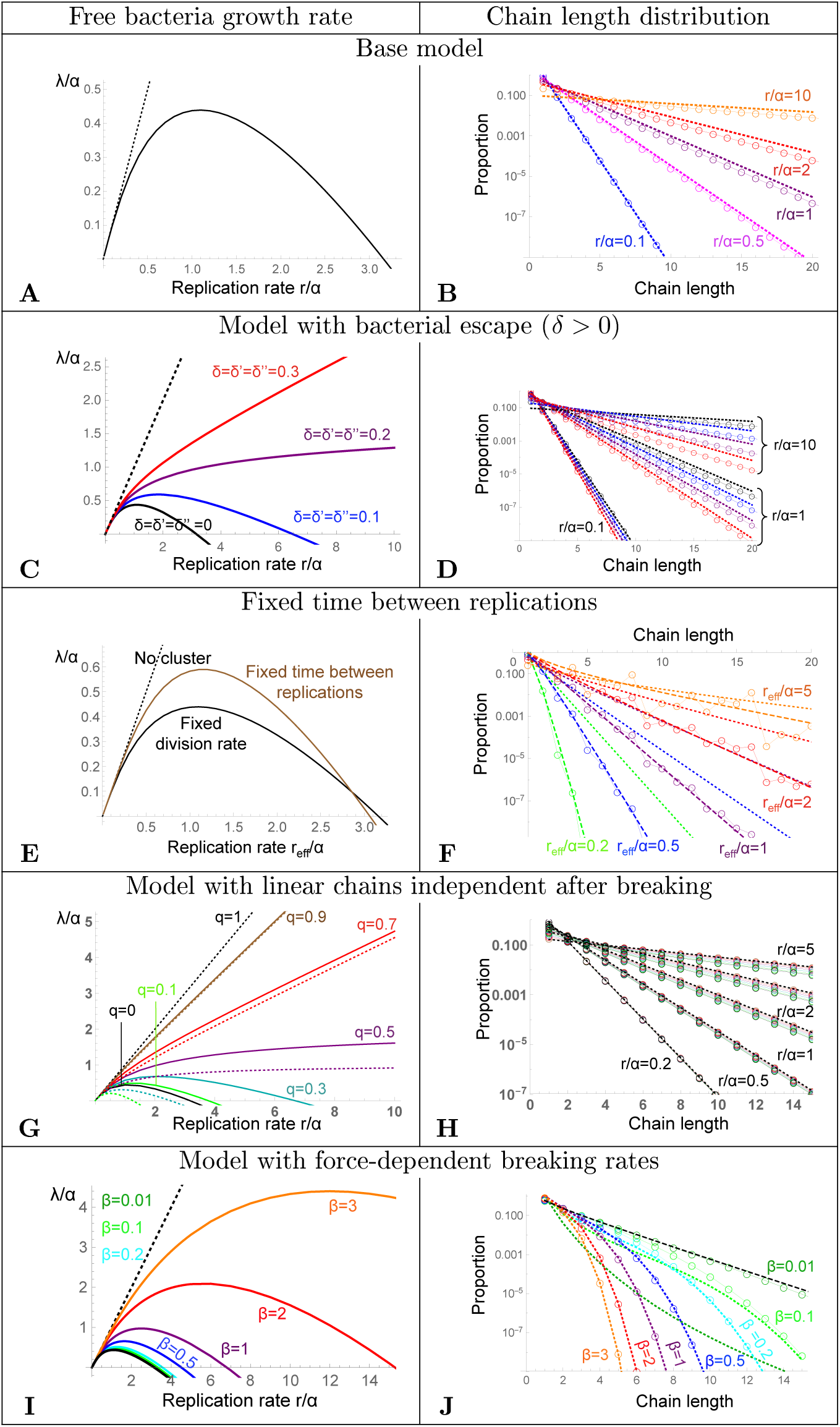
Left panels A, C, E, G, I: Growth rate *λ* of the free bacteria as a function of the bacteria replication rate *r*, both in units of *α*. Numerical results (solid colored lines), and limit with no clusters (*λ* = *r*) (black dotted line). The base model is represented in black in all the left panel figures to ease comparison between models. **Right panels B, D, F, H, J :** Chain length distribution. Open circles linked by solid lines: numerical results. **A, B Base model,** *n*_*max*_ = 40. B. dotted lines: approximation (4) (almost overlaid with the numerical results for *r/α* = 0.1). **C, D Model with bacterial escape.** *δ* = *δ*′ = *δ*″ = 0, 0.1, 0.2, 0.3. *c* = *c*′ = 0, *n*_*max*_ = 40. D. dotted lines: approximation (5). **E, F Fixed time between replications.** *r*_*eff*_ = log(2)*/τ*. *n*_*max*_ = 32. F. approximation (8) (dashed lines), numerical result in the base model (dotted lines). *r*_*eff*_ */α* = 0.2, 0.5, 1, 2, 5. **G, H Model with linear chains independent after breaking**. . G. The dotted black line is the case *q* = 1, for which *λ* = *r*, like in the absence of clusters. The colored dotted lines are the analytical approximation (11). *n*_*max*_ = 200. H. The dotted black lines are the approximate distribution (10) for each *r/α*, which is the exact distribution for *q* = 1. The colours represent the same *q* values than for the left panel. All curves are almost overlaid for small *r*. *n*_*max*_ = 80. **I,J Model with force-dependent breaking rates.** Each color represents a different *β*: *β* = 0.0, (*n*_*max*_ = 20), *β* = 0., (*n*_*max*_ = 15), *β* = 0.2 (*n*_*max*_ = 15), *β* = 0.5 (*n*_*max*_ = 15), *β* =, (*n*_*max*_ = 15), *β* = 2 (*n*_*max*_ = 10), *β* = 3 (*n*_*max*_ = 10). I. The black line is the numerical result for the base model, equivalent to *β* = 0. The curve for *β* = 0.1, (dark green) is almost overlaid with the curve for *β* = 0. J. Chain length distribution for *r/α* = 1. The colored dotted lines the analytical approximation (14), and the black dashed line the numerical result for the base model.

### Chain length distribution

In the long time limit, the number of chains of length *i* is of the order of *Cp*_*i*_ exp(*λt*), with *λ* the largest eigen value. Equation (2) simplifies to:

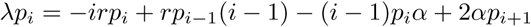

Assuming that *i* is large,

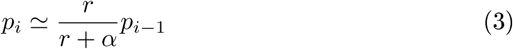

is required. Using this approximation for all *i*, the proportion of chains of length *k* among linear chains and free bacteria is:

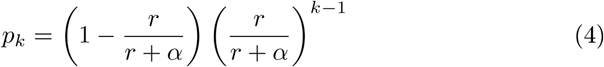

This approximation works relatively well, especially for smaller *r/α* values (see figure 2B). Part of the discrepancy is that equation (3) is an approximation for large *i*, and thus does not hold at small chain length.

## Model with bacteria escape

### Equations

This is similar to the base model presented before, except that we take into account that upon replication, bacteria may not be perfectly bound, and may escape (pannel B of figure 1). We denote *δ* the probability for the two daughter bacteria to become free bacteria upon replication of a free bacterium. We denote *δ*′ the probability that when a bacteria at the tip of a chain replicates, the daughter bacterium on the outside of the chain escapes the enchainment. We denote *δ*″ the probability that when a bacterium at the interior of the chain divides, the daughter bacteria will not be enchained, effectively clipping the chain in two. As free bacteria are more motile than clusters, then *δ* ≥ *δ*′ ≥ *δ*″. We also add here the possibility that the loss rate *c* for free bacteria and *c*′ for chains are different. Then the base equations are:

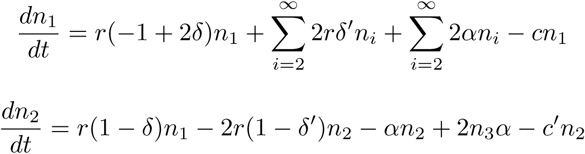

and for *i* ≥ 3,

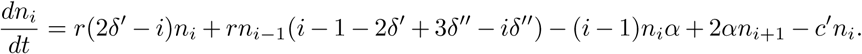

### Free bacteria growth rate as a function of the bacterial replication rate

Similarly to the base model, we study numerically the growth rate as a function of the replication rate (see figure 2C). The larger the replication rate, the more the deviation between the growth rate and the replication rate, which would be its value in the absence of clusters. If *δ, δ*′, *δ*″ are small enough, the qualitative behavior is similar to the base model. But for larger *δ, δ*′ and *δ*″, the growth rate continues to increase monotonically with the replication rate. The same is true when *δ, δ*′ and *δ*″ are different (see supplementary figure 1). If *c* = *c*′, the growth rate is simply offset by minus the loss rate (see supplementary figure 1), and if *c* ≠*c*′, the effect is more complex, but for small *r/α* values it corresponds to an offset of *−c*.

### Chain length distribution

We can reason similarly to the base model (more details in section 3 in appendix), and find:

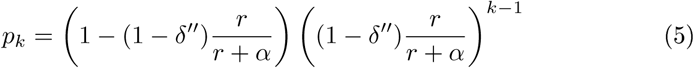

This approximation works relatively well (figure 2D, and supplementary figure 2). The approximation (5) depends on *δ*″, but neither on *δ* nor *δ*′, but *δ* and *δ*′ could actually matter when *i* is small, and indeed we observe (see supplementary figures 3 and 4) that the approximation (5) works slightly less well when *δ*″ is different from *δ* or *δ*′. If *c* = *c*′, the distribution is exactly the same as for *c* = *c*′ = 0, and if *c* ≠*c*′, the distribution changes very little (see supplementary figure 5).

**Figure 3:**
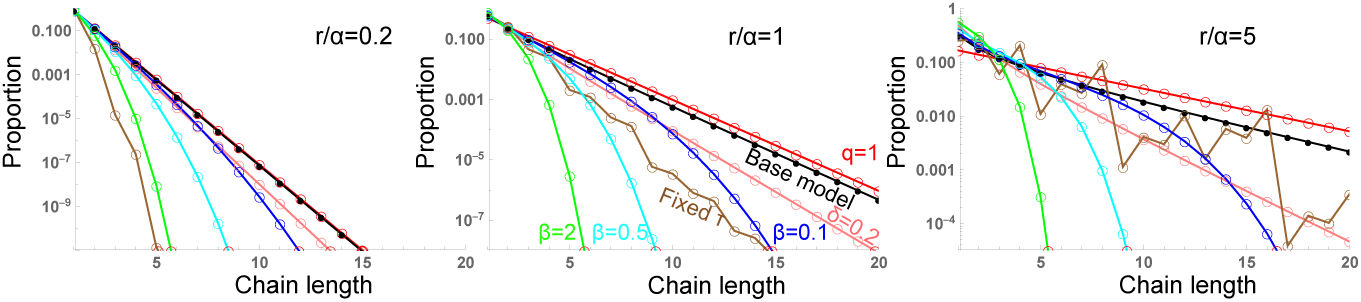
Comparison of the chain length distributions for the different models. The base model is represented in black. For the case with bacterial escape when a bacterium replicates, one numerical value is represented (*δ* = *δ*′ = *δ*″) (pink), the lower this value, the closer to the base model. The model with fixed replication time is represented in brown (for this model we choose *τ* = *log*(2)*/r*). The model with linear chains independent after breaking (*q* > 0) is shown in red for *q* = 1, the most different from the base model. All intermediate *q* values are between the black and the red curves. The model with the force dependent breaking rate is represented for 3 values of *β*: 0., (blue), 0.5 (cyan), 2 (green). All the results are numerical, using *n*_*max*_ values as in figure 2, except for *q* = 1 which is an exact analytical result.

**Figure 4:**
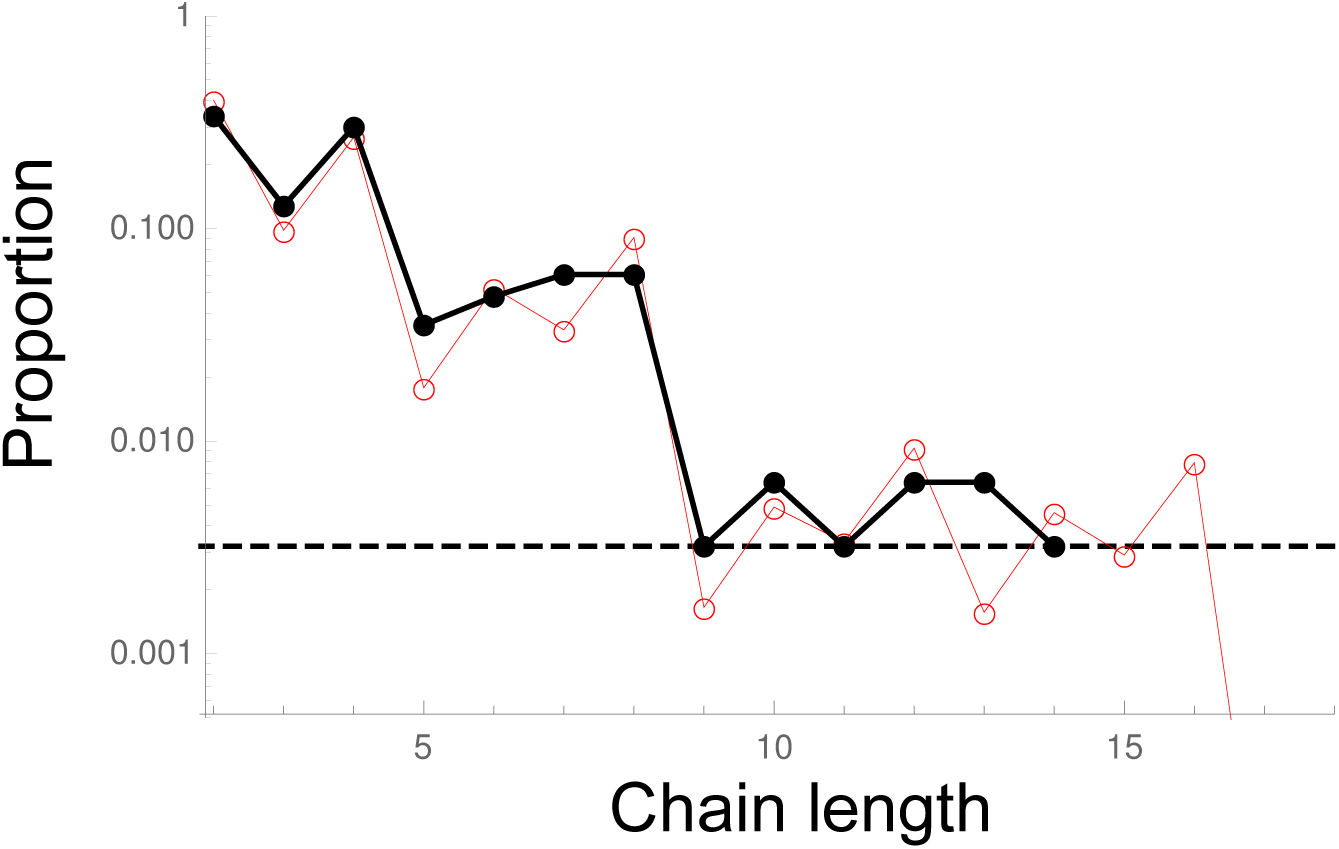
Comparison with experiments. Chain length distribution (proportions relative to the linear chains of length ≥ 2 are represented in log scale). The black dots and line are the experimental data. The horizontal dotted black line represents the case in which there is one chain of the given size. No chains longer than size 14 were detected in our experimental images. The red line and points are the numerical results for the model with fixed replication time. In this model, there is only one free parameter, *r*_*eff*_ */α* = log(2)*/*(*ατ)*, which fitted value is 4.1(see appendix 7 for more details).

### Model with fixed replication time

In this variant of the base model, bacteria divide every *τ*. The effective growth rate is *r*_*eff*_ such that exp(*r*_*eff*_ *t*) = 2^*t/τ*^, thus *r*_*eff*_ = log(2)*/τ*.

#### Equations

Let us start by considering a chain of *n* bacteria at *t* = 0, right after a replication event. Let us denote *l*(*n, i, t*) the probability that at *t*, this chain has lost *i* bacteria in total on the extremities, and consequently is of length *n* − *i* at *t* (*n* ≥ 2 and 0 ≤ *i* ≤ *n*–2). Before the next replication event, since we assume *q* = 0 as in the base model (meaning that if the chain breaks somewhere else, the subparts form a more complex cluster and thus are “lost” for the system), we have:

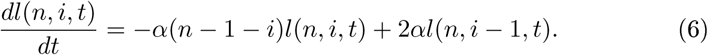

At *t* = 0, *l*(*n*, 0, 0) =1 and for 0 < *i* < *n–*1, *l*(*n, i*, 0) = 0. It can be checked easily that the following expression is the solution of (6), with *l*(*n*, 0, 0) = 1 and *l*(*n, i* > 0, 0) = 0, for any 0 ≤ *i* ≤ *n* − 2:

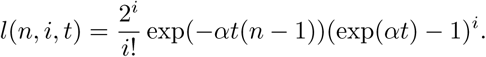

Any chain of length > 2, has two outermost links, each breaking at rate *α*, liberating one free bacterium. A chain of length 2 breaks at rate *α*, but liberates two free bacteria. Starting from a unique chain of length *n*, the probability that a linear chain of length *n − i* is present at time *t* is *l*(*n, i, t*). Consequently, the total rate of production of free bacteria at time 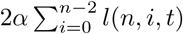. Thus, the average number of free bacteria generated during *τ* by this chain of *n* bacteria is:

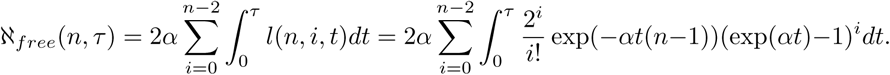

A chain of length *n* right before replication becomes a chain of length 2*n* upon it, and will have contributed to chains of length *k* by *l*(2*n*, 2*n − k, τ)*, and to free bacteria by ℵ_*free*_(2*n, τ)* right before the next replication event. With *n*_*i*_(*t*) the number of chains of length *i* at time *t* (right before a replication event), we can write the matricial relation between the *n*_*i*_(*t*) and *n*_*i*_(*t* + *τ)* (right before the next replication event) as follows:

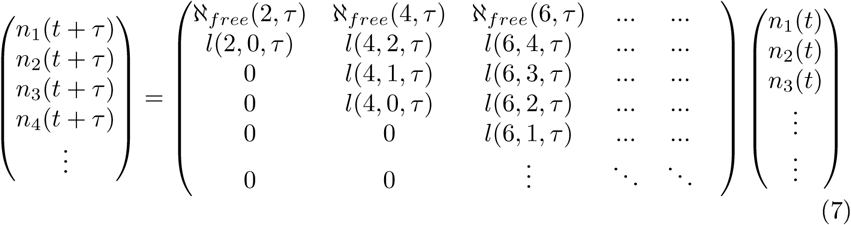

This matrix is then cut to size *n*_*max*_×*n*_*max*_, and the corresponding eigen-system is solved numerically.

#### Free bacteria growth rate as a function of the bacterial replication rate

The shape of the relation between free bacteria growth rate and (effective) replication rate (figure 2E) is very similar in the fixed replication time vs. fixed replication rate models, with a maximum of the growth rate for a finite value of the (effective) replication rate, at close values (*r*_*eff*_ = 1.15*α* vs. *r* = 1.09*α*). When the replication is at fixed time intervals instead of a fixed replication rate, the maximum growth rate is higher, and it dips faster at increasing effective replication rate. Indeed, in the case of fixed replication rate, the distribution of durations between two replications is exponential, thus more spread. Close to the maximum, the presence of short replication intervals makes that there can be more cluster formation, and conversely, at higher replication rates, the presence of longer replication intervals results in more production of free bacteria.

#### Chain length distribution

We show here the main steps to calculate analytically an approximation for the chain length distribution, and more details are given in section 4 in appendix. We define *n*_*i*_(*t*) the number of chains of length *i* at *t* with *t* taken just before a replication. Assuming *i* even,

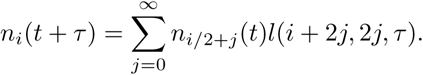

In the long time limit, *n*_*i*_(*t*) = *Cp*_*i*_ exp(*λt*), with *λ* the long term growth rate, that is such that exp(*λτ)* = 𝒩, with 𝒩 the largest eigenvalue of the matrix of equation 7. Then previous equation leads to:

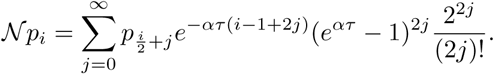

We make the assumption that the first term of the sum is large compared to the rest of the sum (assumption discussed in appendix, section 4). Then,

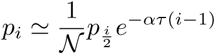

and recursively, for *i* = 2^*k*^, with *k* integer,

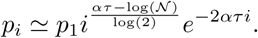

When *ατ* ≫1, links typically break before the next replication1thus there is little impact of the clustering on the growth. Consequently, the growth will be close to its valuein theabsenceof clustering1i.e. doublingevery *τ*, and thusin thislimit 𝒩= 2:

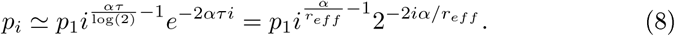

This rough approximation allows to explain the core of the observed distribution (figure 2F). There are bumps, due to the replication every *τ* (which in the absence of link breaking would results in chains of length 2^*k*^ only), which makes that chains of power-of-two length are overrepresented. Compared to the case with fixed replication rate, the distribution is much narrower.

### Model with linear chains independent after breaking (*q* > 0)

#### Limit case: subchains always remain independent linear chains after breaking (*q* = 1)

In this model, when a chain breaks, the two resulting chains remain independent and can thus continue to participate in the dynamics of the system:

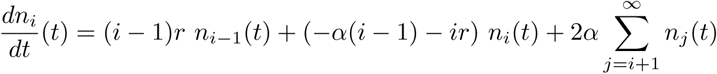

We recognize here the equation studied in[18], where they described chains of growing unicellular algae. As it has been shown, the steady state solution of the system is:

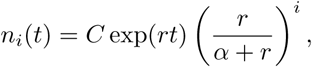

with *C* a constant dependent on the initial state of the system. In the steady state, the growth rate is equal to the replication rate. Note that the resulting chain length distribution is then exactly equal to its approximation in the base model (4). The average chain length is 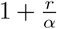, which shows that, as expected, if the link breaking rate is high compared to the replication rate (*r/α*≪1), the average length is close to one as no cluster has the time to form: all the bacteria remain free.

### Intermediate case: chains can either be independent or trapped after breaking

More realistically, after breaking, chains will have some probability to either encounter each other and remain trapped in more complex clusters, or to escape and become independent. We will assume in the following that if a chain of length *N* breaks at a link at the extremity, releasing a chain of length *N*, and a free bacterium, then the free bacterium, smaller and likely more mobile, will escape in all cases; but that if the link that breaks is elsewhere, the probability for the new chains of lengths *N − k* and *k* (*k* > 1) to escape and continue as two independent linear chains will be *q*, and the probability that they bind and form a more complex cluster will be 1*- q*, with *q* independent of *k*. We write the equations for the number *n*_*i*_(*t*) of chain of *i* bacteria:

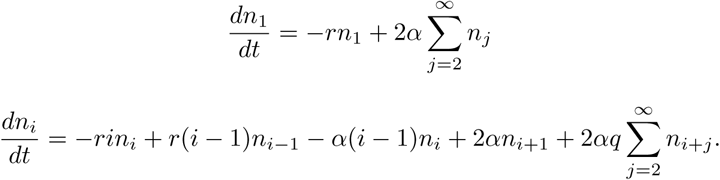

In the long time, *n*_*i*_(*t*) *→ Cp*_*i*_ exp(*λt*) with *λ* the largest eigenvalue.

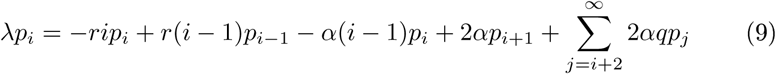

This is valid for any *i*. We assume that *p*_*i*_ decreases fast enough with *i* such that the sum from *i* + 2 to ∞ of the *p*_*i*_ is an order of magnitude less than *ip*_*i*_. Then, the largest elements of equation (9) when *i* is large enough are the terms multiplied by *i*, and consequently:

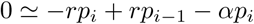

Leading to

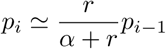

If this is valid for any *i*, theproportion of chains of length *k* among linear chains and free bacteria is:

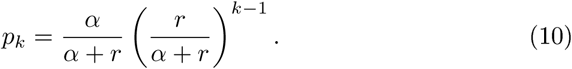

Note that this expression does not depend on *q* and is the same as the approximation for the base model (4) and the exact expression for *q* = 1. We compare this approximation with the numerical results and they are in good agreement (figure 2H), except when both *q* is small and *r/α* is large, and even in this case it gives a reasonable approximation.

Replacing *p*_*i*_ by its expression (10), equation (9) simplifies to:

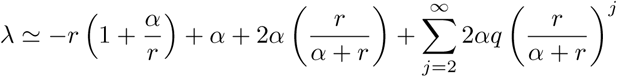

which after simplifications leads to:

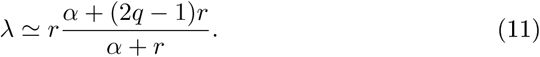

This approximation does not work for *q* < 0.5, but it works well for *q* close to 1, and gives the right slope for *r/α* large for *q* > 0.5 (figure 2G). We can observe that for *q* > 0.5, *λ* increases indefinitely when *r* increases; but it has an intermediate maximum for *q* < 0.5. Intuitively, if *q* > 0.5, when a chain breaks it leads to more than one independent linear chain, thus the population of linear chains and thus free bacteria may increase monotonically with *r/α*, whereas if *q* < 0.5, chains that break lead to less than one independent chain on average, and thus the behaviour of the system is more determined by the fate of the chains, and thus closer to the results for *q* = 0, for which the growth rate *λ* has a maximum as a function of *r*.

The proportion of free bacteria relative to the total number of bacteria is proportional to exp(*λt*)*/* exp(*rt*), which, using approximation (11) tends to exp(−2*r*^2^*t*(1−*q*)*/*(*r* + *α*)). The proportion of free bacteria thus decreases over time, and decreases with increasing replication rate *r*. Thus the proportion of bacteria trapped in clusters, which is 1 minus the proportion of free bacteria, increases with increasing replication rate even when *q* > 0.5.

### Model with force-dependent breaking rate

#### Equations

What drives link breakage? The links could break if there was some process degrading the sIgA, but the sIgA are thought to be very stable[19]. Another possible explanation for link breaking is that the bound antigen can be extracted from the bacterial membrane, at a rate which may vary exponentially with the force[20][15]. The forces applied on the links are likely mostly due to the hydrodynamic forces exerted by the digesta flow on the bacterial chain. Taking the linear chain as a string of beads, as done for polymer chains, and in a flow with a constant shear rate, the force is predicted to be larger as the chain grows longer, and the largest at the center of the chain[17]. A more detailed discussion and the calculations can be found in section 5.1 in the appendix. Taking *α* as the breaking rate in the absence of shear, and *β* a constant expressing the strength of the coupling between hydrodynamic forces and link breaking, the resulting equations for this minimal model taking into account the forces are:

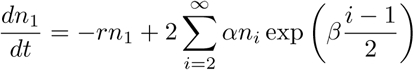

and for *i* even,

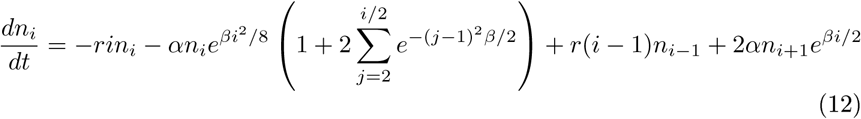

and for *i* > 1 odd,

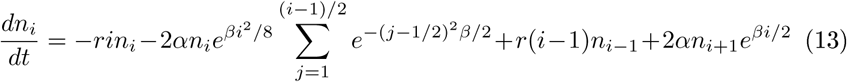

### Free bacteria growth rate as a function of the bacterial replication rate

The growth rate as a function of the replication rate has a qualitatively similar shape as for the base model (figure 2I), with a finite replication rate maximizing the growth rate. The limit *β*→0 corresponds well to the base model, as expected. When *β* increases, the replication rate maximizing the growth rate increases, as the effective breaking rate is higher. Numerically, we find (see supplementary figure 7) that the replication rate maximizing the growth rate scales as *α* exp(0.8*β*).

### Chain length distribution

Similarly to the other models, for *t* long enough, *n*_*i*_ *≃ Cp*_*i*_ exp(*λt*) (with *λ* the largest eigenvalue), and assessing which terms in equations (12) and (13) will be dominant, we ultimately obtain (details in supplementary section 5.3):

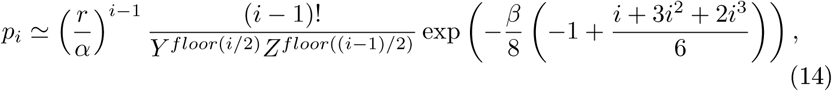

with 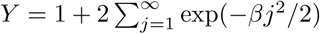 and 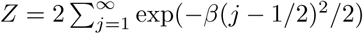. This approximation works well, except for small *β* (figure 2J, and supplementary figure 8). Compared to the base model, the number of chains decreases much faster with their length. Indeed, the breaking rates for each link increase importantly with the chain length, thus larger chains are much less stable than in the base model.

## Comparison with experimental data

We analyzed (see section 7 in the appendix) microscopy images of cecal content from vaccinated mice infected with *S*.Typhimurium, which were acquired for our previous study [12]. Most clusters are large, and of complex shape. But smaller clusters are linear, and we obtained the distribution shown by the black line and points in figure 4. The model with fixed division time is *a priori* more realistic. The best fit is obtained with this model, with the one adjustable parameter *r/α* = 4.1 (red line and points). It seems however, though there is not enough data to quantitatively ensure it, that there are less long clusters than expected (see appendix section 7.3 for an expanded discussion). The data may be biased, as longer chains may not be fully in the focal plane. That the distribution is relatively narrow could also be compatible with force-dependent breaking rates.

## Summary of results and discussion

We started from the recent finding[12] that the protection effect of sIgA, the main effector of the adaptive immune system in the gut, can be explained by enchained growth. Because sIgA are multivalent, they can stick identical bacteria together if they encounter each other. Early in infection, bacteria of the same type are at low density, thus typical encounter times are very long, but when a bacterium replicates, the daughter bacteria are in contact and thus can remain enchained to each other by IgA. Bacteria in clusters are less motile than individual bacteria, and in particular, are not observed close to the epithelial cells. In the case of wild type *S*.Typhimurium, only free bacteria which can interact with the epithelial cells contribute to the next steps of the infection process. Despite the presence of sIgA, some free bacteria are observed. It could be that they escape at the moment of replication. But, along with the observation that clusters do not grow indefinitely, it could also be a sign that the links between bacteria break. It is also physically expected that the links have some finite breaking rate. If the typical time between two bacterial divisions is much larger than the typical time for the link to break, then there would be no cluster. Conversely, in the inverse case, bacteria will be very likely to be trapped in large clusters. Then, even if sIgA are produced against all bacterial types, the bacteria dividing faster will be disproportionately affected.

We investigated if this qualitative idea holds with more realistic models. We started from a base model in which: bacteria replicate at a fixed rate; remain enchained upon replication; until the link between them breaks at a given fixed breaking rate, identical for all links; and considering that, because of the way bacteria such as *Salmonella* or *E.coli* divide, the early clusters are linear chains of bacteria; when the chain breaks at an outermost link, we assumed the free bacteria will escape; but if the chain breaks elsewhere, we assumed that the two resulting sub-chains encounter each other quickly and form clusters of more complex shapes from which individual bacteria do not escape. We studied this base model with a combination of analytical and numerical approaches. We also tested the robustness of our findings by studying separately several variations of the base model: a probability of escaping upon replication, loss rates, fixed replication time, non-zero probability for the subchains to escape, and force-dependent breaking-rates. For each model, we studied how the growth rate of the free bacteria varies with the replication rate (which would be equal if there were no clusters), and the distribution of chain lengths.

We find that, except in the very specific case in which subchains always escape upon link (*q* = 1), the growth rate of the number of free bacteria is lower than the replication rate. And more spectacularly, in most of the models studied (but not if more than half the subchains escape upon link breaking, or if there is a significant probability for bacteria to escape enchainement upon replication), the growth rate of the number of free bacteria is non-monotonic with the replication rate : there is a finite replication rate which maximizes the growth rate of non-clustered bacteria. At very high replication rates, bacteria get trapped in more complex clusters and cannot contribute anymore to the free bacteria dynamics and thus to the next steps of the infection process. The replication rate maximizing the growth rate is of the order of the breaking rate, though its specific value depends on the details of the model. To summarize, except when *q* = 1, we always find that the higher the replication rate, the higher the proportion of bacteria trapped into clusters; and in many cases, the effect is even more dramatic, with the growth rate of free bacteria that may decrease with the replication rate.

The chain length distribution is dependent on the model (see figure 3). In most cases, the proportion of linear chains having length *k* decreases as *γ*^*k*^, with *γ* some constant smaller than 1. When replication occurs at fixed time, or when breaking rates are force-dependent, the proportion of longer chains decreases faster. There are models with different chain length distributions but qualitatively similar dependence of the growth rate on the replication rate, and the opposite is true too. This shows that large clusters have little importance for free bacteria production, what matters most is the small chains dynamics. It is reassuring, as we did not consider buckling, which would make long linear chains fold on themselves and produce more complex clusters, and may bias the linear chain distribution for very large lengths. It should also be noted that with fixed division time, not only the distribution is bumpy, as chains comprising a power of two number of bacteria are more frequent than others, but the distribution is also narrower.

We analyzed experimental data on clusters of *S*.Typhimurium in the cecum of vaccinated mice. The experimental chain length distribution is in line with the model of fixed replication time, which is indeed more realistic. There is however somewhat less large chains than expected. More data would be necessary to asses this more reliably. This could be because of possible bias in the data. This could be also compatible with force-dependent breaking rates. Additional experiments, for instance to measure the breaking rate, could help by giving additional independent information and constrain the fitting. To test the dependence of the growth rate with the replication rate, an ideal experiment would be to compare similar bacterial strains, but with differing replication rates, and compete them in the same individual. It is however very challenging to obtain bacteria that differ only by their replication rate, particularly in vivo.

sIgA-enchained bacterial clusters could be studied in vitro to measure how they break. However, using in vitro results to draw conclusions on in vivo systems is limited. First, there could be chemical or enzymatic components of the lumen that could facilitate or hinder link breaking, and the non-Newtonian viscosity of the digesta could play a role in the mechanic forces felt by the links, thus a simple buffer may not mimic well the real conditions. More crucially, the exact forces felt by particles of the size of bacterial clusters are not well characterized. Most studies of the flow characteristics in the digestive system rely either on external observations of the peristaltic muscles[21] or indirect measures of times for a marker to exit some section of the digestive track[22]. More quantitative study of the digestive flow at small scales is just beginning[23, 24, 25, 26, 27, 9, 10] and in the future it may give more clues to assess to which forces bacteria are subjected to in the digestive track.

The mechanism we propose is nevertheless plausible. The observation in vaccinated mice of the existence of single bacteria and small clusters, and particularly small linear chains with an odd number of bacteria, are pieces of evidence that clusters do break in these in vivo conditions. An alternative explanation could be that some bacteria escape enchainement upon replication. However, at higher bacterial densities, we have evidence of independent bacteria binding when they encounter[12], thus sIgA coated bacteria are adhesive. When two daughter bacteria divide, they are in contact, thus if sIgA is adhesive, escape is unlikely (see appendix section 2). Importantly, even though our results show that specific conditions are needed for the growth rate to decrease with high replication rates, we almost always find that the higher the replication rate, the higher the proportion of bacteria trapped in clusters. Thus, even when it does not reverse the relationship between the growth rate of the free bacteria and the replication rate, it is at least dampening this relationship, and can be a tool both to control pathogenic bacteria, but also to maintain homeostasis of the gut microbiota. It is also interesting that there are other host effectors besides sIgA that bind bacteria together: neutrophil extracellular traps for instance[28] and there could also be an interplay between replication rates and the breaking of the links mediated by these other effectors, as the mechanism we propose here is generic.

As for any mechanism to fight against bacteria, how easily resistance can be evolved is crucial. On the one hand, the replication rate could evolve. But bacteria replicating slower would be less competitive with other bacteria in the absence of sIgA, and a slower growth leaves more time for further host response. On the other hand the typical link breaking time could evolve. On the host side, sIgA is thought to be mechanically very stable, and experiments about the bonding of cells by sIgA seem to point to the link failing because of the extraction of the antigen rather than because of sIgA breaking, and rather than the sIgA/antigen bond detaching[20][14]. In the case of IgA defficiency, there is more secretion of IgM, and microbiota is disturbed[29]: we may speculate that IgM being less powerful for microbiota homeostasis is related to these immunoglobulins being more protease-sensitive than IgA and thus cleaved on shorter time scales[30]. On the other side, bacteria could evolve surface antigens. It could be interesting to think that bacteria could produce decoy antigens with no functional value, but against which the immune system will mount an immune response, and that are more easily released from the bacteria, thus disabling the main sIgA mode of action (being easily evolvable would also be a benefit). Such decoys would however be a metabolic cost for the bacteria, and when breaking, may unmask other antigens corresponding to crucial functions of the bacteria. It could be argued that the capsule around bacteria such as *Salmonella* spp., and also common in pathogenic *E.coli*, may behave as a decoy, though it has also other functions. Evolving resistance to IgA-mediated enchainment would thus be costly.

Along the same lines, we may speculate whether mechanical aspects could be a reason why sIgA against some antigens are not efficient for protection. For instance, while anti-flagella sIgA aggregate very well *Salmonella* Enteriditis together, they are not efficient for protection[31]. A main reason could be that as *Salmonella* can switch flagella production on and off, then some *Salmonella* will always escape these sIgA, and seed the infection[32]. An additional possibility could be that flagella may more easily break, especially as distance between bacteria bound by flagella (long) is likely larger than for bacteria bound by O-antigens (on chains shorter than flagellas)[33], and thus the shear forces would be larger. Further, the mechanical properties of the outer sugar layer of the gram negative bacteria could vary, and thus could be used to tune interactions. However, it would add another constraint on bacteria, and the general result that the growth rate compared to the replication rate is at least dampened by the cluster formation would remain.

In the crowded environment of the gut, it is hard for the host to identify the good and the bad bacteria. That vaccination with dead bacteria is sufficient to produce sIgA and protection, shows that the host does not discriminate well against which bacteria they produce sIgA, as these dead bacteria do not harm. Linking the effect (here the clustering) of the immune effectors with a property directly relevant to the potential bacterial pathogeneicity (here the replication rate) avoids to make complex decisions about which bacteria to produce effectors against.

## Acknowledgments

The authors thank Anne-Florence Bitbol for careful reading of an earlier version of the manuscript, as well as other Laboratoire Jean Perrin members for useful discussions, in particular Raphaël Voituriez. We also thank Roger Lentle for giving useful indications about the flow in the mouse digestive system.

## Appendix

### 1 Order of magnitude of the encounter time between two bacteria

The typical time to find one target of radius *a* in a sphere of radius *b* by diffusion is of the order of *b*^3^*/*(*Da*), so the typical time when there are *N* bacteria in a volume *V* is of the order of *V/*(*NDa*). For bacteria, *a* is in the micrometer range. Bacteria such as salmonella or E.coli typically swim at 10*μm/s*, and change direction every second, which gives a diffusion coefficient of the order of 10 ^− 10^*m*^2^*/s* [1, 2, 3] (The peristaltic motions of the digesta are large scale movement rather than local diffusion, so we assume they have a smaller effect on diffusion). The mouse’s cecum has a volume of the order of (1cm) ^3^. In experiments of [4], the smallest inoculum consists in *N* = 10^5^ bacteria, which is already large compared to what could be a realistic number of pathogenic bacteria in food poisoning (10^5^ is the typical number of Salmonella for food poisoning in humans [5], which are much larger than mice). With these numbers, the typical encounter time is of the order of 10^5^*s*, i.e 30h, about 10 times longer than the typical digestion time in mice.

### 2 Argument for high enchainment probability upon replication

When a bacterium replicates, the time for septation is of the order of a few minutes. We intuitively think that this time is much larger than the time *τ*_*k*_ required for bacteria to stick together when they randomly meet. The aim of this section is to check this intuition by giving an overestimate of *τ*_*k*_.

If the diffusion coefficient is high enough, the time for bacteria to stick to each other will be limited by which proportion of the time they spend in close vicinity, and the rate *k* at which bacteria stick to each other when they are in close vicinity, *k* being the inverse of *τ*_*k*_. If the diffusion coefficient is smaller, then the time to first encounter will also play a role, but as we calculate an overestimate of *τ*_*k*_, we can neglect this scenario.

We use the data on figure 1k of [4] about non-dividing bacteria (so the only sticking is from random encounters). The majority of them are aggregated after *τ*_*exp*_ up to 8 hours (from the inoculum ingestion to the sampling used for imaging) for a concentration of 10^7^ *−* 10^8^ bacteria. As we will see, this estimate of *τ*_*k*_ is proportional to *τ*_*exp*_ and *N*, so to be conservative, as we will calculate an overestimate of *τ*_*k*_, we take the highest concentration and the maximum experimental time, i.e. *N* = 10^8^ bacteria in *V* = 1*cm*^3^ (cecum volume) and *τ*_*exp*_ = 8 hour.

The bacteria typical size is a few micrometers, we thus take 3*μm* as an overestimate of the maximum bacterial size. Thus to be in close contact, two bacteria must be at most at *a* = 3*μm* away. Let us assume that then, the volume of possible contact is 4*/*3*πa*^3^, which is also an overestimate, because only certain orientations will allow bacteria to touch each other. Then, the proportion of time spent in close contact will be of the order of (*N* 4*πa*^3^)*/*(3*V)*.Then the typical time to stick to each other will be *τ*_*exp*_ = *τ*_*k*_3*V /*(*N* 4*πa*^3^). Then *τ*_*k*_ = *τ*_*exp*_*N* 4*πa*^3^*/*(3*V)*. Numerically, we obtain about 5 minutes as an overestimate of *τ*_*k*_.

Note that this is a large overestimate. Indeed, when bacteria get clumped to each other, their effective concentration decreases, thus it takes longer for the last bacteria to meet others, and thus the time for most bacteria to be clumped will be significantly larger than the inverse of the early clumping rate.

With all these highly conservative estimates, we find *τ*_*k*_ at the very most of the same order of magnitude as the septation time, and very likely much smaller. Hence the probability for bacteria to escape enchainment is small, which justifies that we take in general the limit of no escape.

### 3 Model with bacterial escape (*δ* > 0) and differential loss (*c* ≠ *c′*)

**Figure S1:**
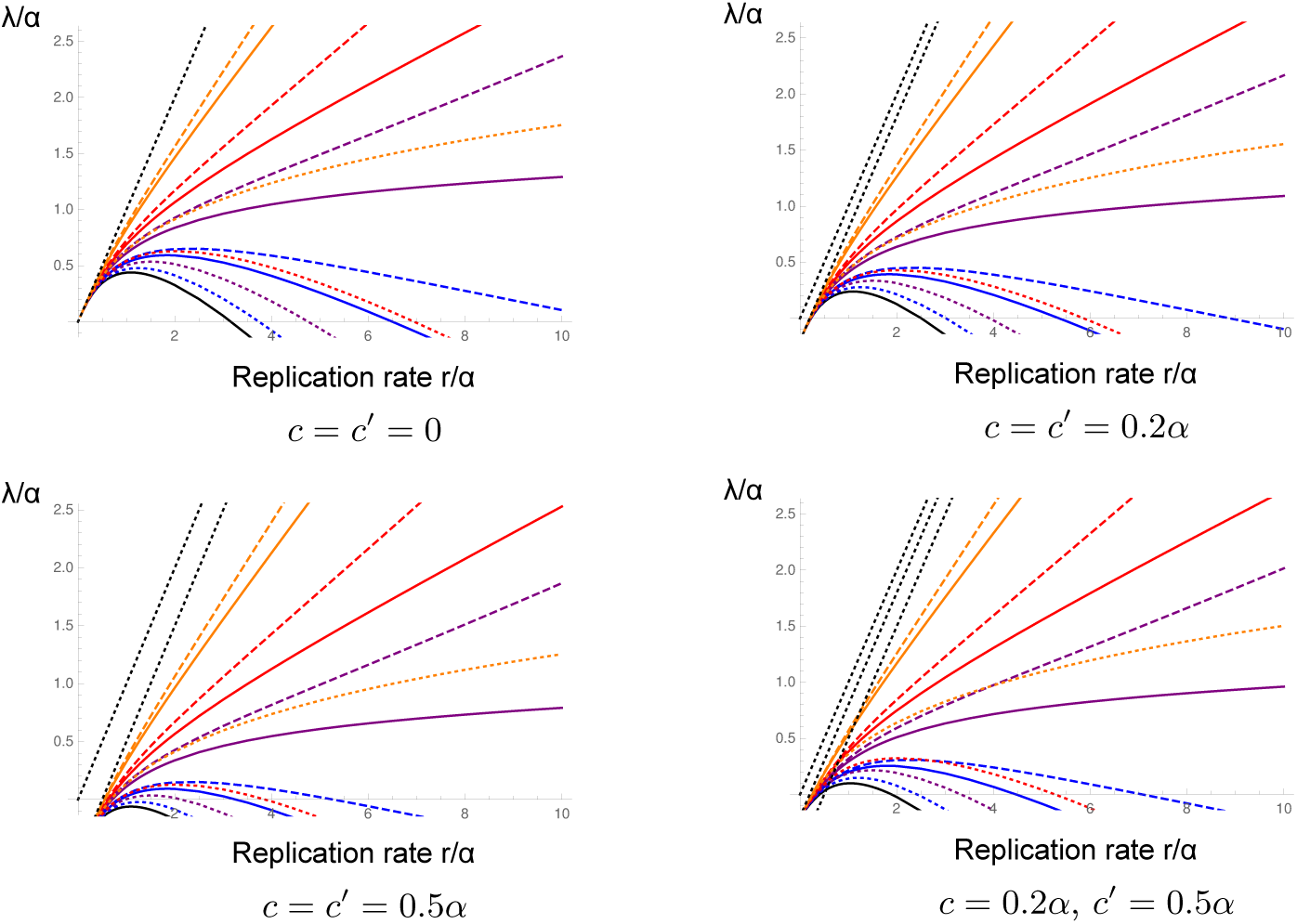
Growth rate *λ* as a function of the replication rate *r*, both in units of *α*. Numerical results (colors), with *δ* = *δ′* = *δ″* (solid lines), *δ* = *δ′*, and *δ″* = 0 (dashed lines), *δ′* = *δ″* = 0 (dotted lines). *δ* = 0, 0.1, 0.2, 0.3, 0.5. Note that for *δ* = 0, solid, dashed and dotted lines collapse, as expected. The black dotted lines are either *r/α*, (*r − c*)*/α* or (*r − c′*)*/α*. As expected, if *c* = *c′*, the resulting growth rate are the same than when *c* = *c′* = 0, minus *c*. If *c* ≠ *c′*, the results are closer for small *r/α* to the results if both *c* and *c′* had the *c* value. For the numerical results, *n*_*max*_ = 40.

Figure S1 shows how the growth rate depends on *r* for different *δ, δ′, δ″, c* and *c′*.

Our numerical study of the system showed us that there is some critical value *δ*_*c*_ below which the behavior is qualitatively similar to the behavior of the system with *δ* = 0, i.e. with a finite maximum of the growth rate of the free bacteria as a function of the replication rate; and above which the growth rate continues to increase with replication rate. Actually, for *δ* > 0.5, the growth rate necessarily continues to increase with the replication rate. Indeed, upon replication, one free bacteria becomes two daughter bacteria, an average of 2*δ* of them staying free. Thus the net gain in free bacteria is 2*δ* − 1. Thus for *δ* > 0.5, the growth rate of free bacteria is at minimum *r*(2*δ* − 1). Consequently, *δ*_*c*_ ≤ 0.5.

We detail here how to obtain the approximation for the chain length distribution. In the long time limit, the number of chains of length *i* is of the order of *Cp*_*i*_ exp(*λt*), with *λ* the largest eigenvalue. Equation (8) of main text simplifies to:

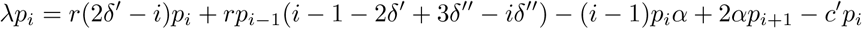

Assuming that *i* is large,

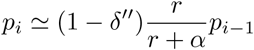

is required. Using this approximation for all *i*, the proportion of chains of length *k* is:

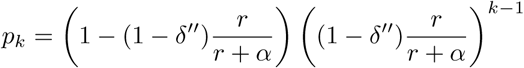

Free bacteria are released at a rate 2*rδ′* + 2*α* per chain. This rate is independent of the chain length. The direct contributions to the increase of free bacteria from chains of length *i* compared to all the larger chains will be (with *K* = (1 *− δ″*)*r/*(*r* + *α*)):

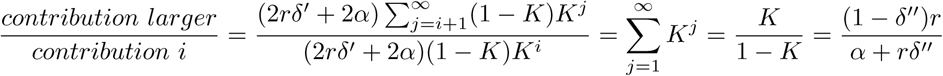

If *r* is small compared to *α* (replication rate ≪ breaking rate), then this ratio is small. Thus the larger chains are quickly negligible. Indeed, in this regime, chains typically dislocate before new replications, so there are few larger chains.

Figures S2, S3, S4, S5 show how the chain length distribution depends on *δ*, *δ′, δ″, c* and *c′*.

**Figure S2:**
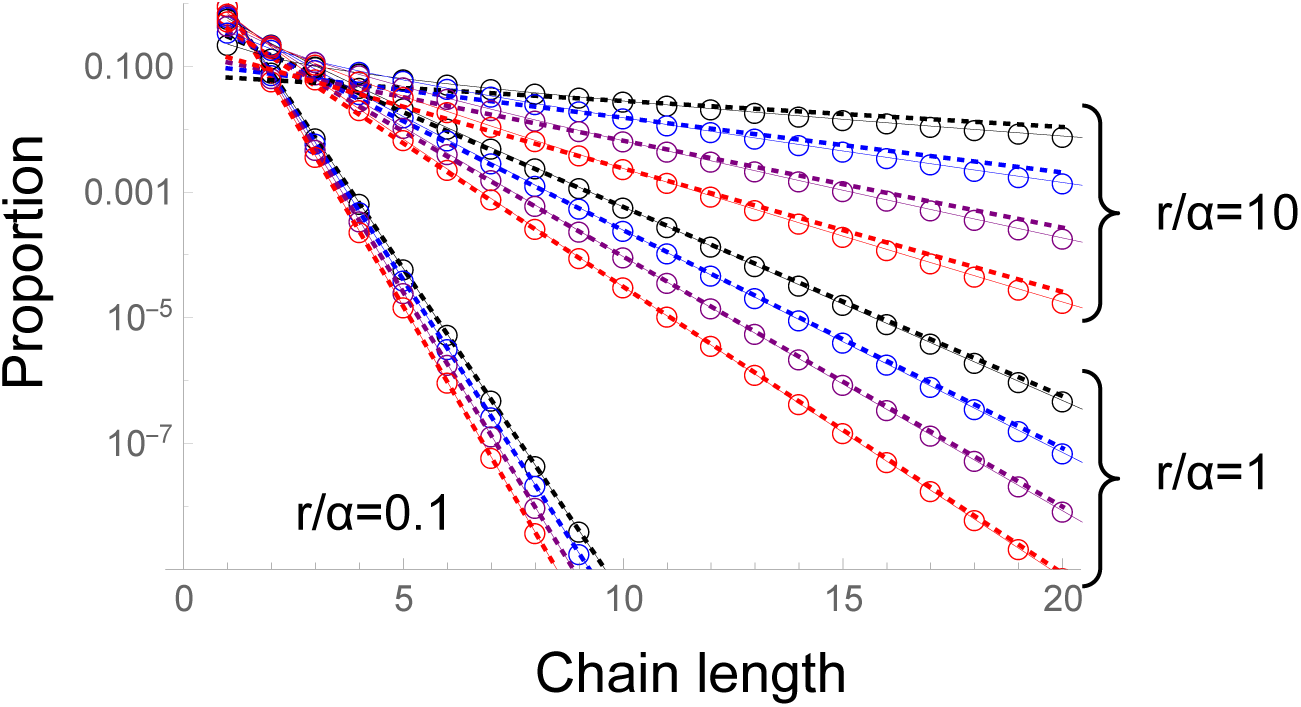
Chain length distribution. All as in main figure 2D (model with bacterial escape) (numerical results: points linked by solid lines), except that the approximation (9) of main text (dotted lines) is rescaled by the numerical value at *n* = 10, i.e. instead of representing log(*p*_*i,approx*_), what is represented is log(*p*_*i,approx*_ *\* p*_10,*numeric*_*/p*_10,*approx*_) = log(*p*_*i,approx*_)+ log(*p*_10,*numeric*_*/p*_10,*approx*_). This shows that the approximation captures well the length distribution of large chains. We do this because the base for the analytical approximation is the ratio *p*_*i*+1_*/p*_*i*_, for which we get a limit expression valid for large *i*. To calculate the whole distribution *p*_*i*_, we assume that this limit expression for the ratio is valid for any *i*, whereas this will not be the case for small *i*. If the limit expression for *p*_*i*+1_*/p*_*i*_ is correct for large *i* but not for small *i*, the slope in log scale plot will be correct, but with some offset dependent on how wrong we got the small *i* case. Making this renormalization enables to check more easily whether the slope is correct. *δ* = *δ′* = *δ″* = 0 (black), 0., (blue), 0.2 (purple), 0.3 (red). *n*_*max*_ = 40.

**Figure S3:**
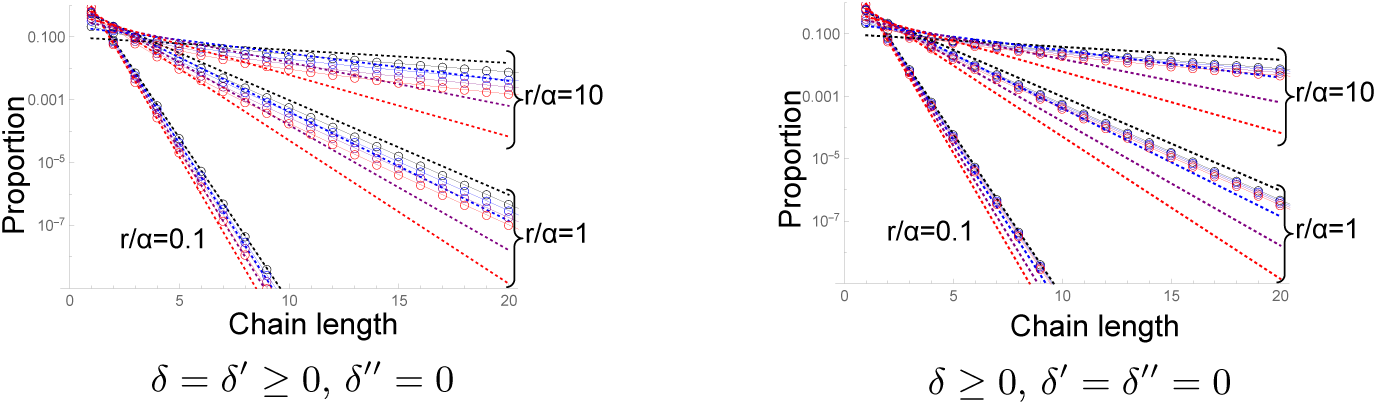
Chain length distribution. All as in figure 2D (model with bacterial escape) (numerical results: points linked by solid lines), except the values of *δ′* and *δ″*: *δ* = 0, 0.1, 0.2, 0.3. Dotted lines: approximation (5) of main text. Approximation (9) of the main text predicts that the distribution should depend only on *δ″*, and not *δ* nor *δ′*. In these figure where *δ″* = 0 but *δ* (and in the left pannel *δ′*) have non-zero values, we do observe that the distribution, in particular its slope, is closest to the result for *δ* = *δ′* = *δ″* = 0. *c* = *c′* = 0, *n*_*max*=40_.

**Figure S4:**
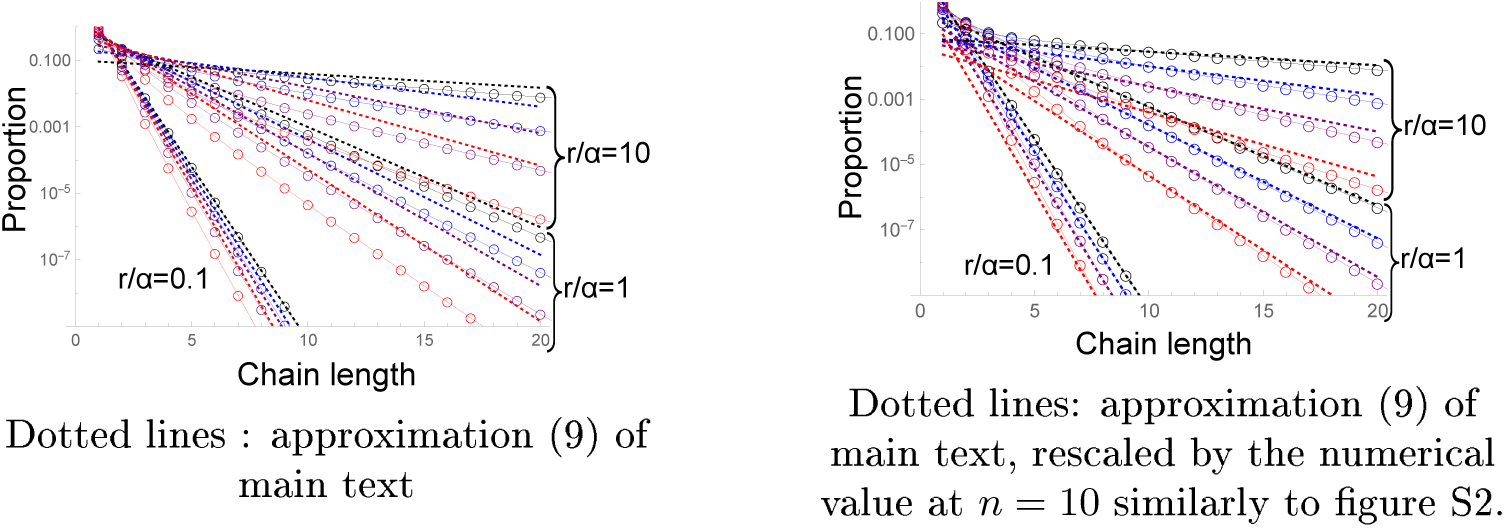
Chain length distribution, for *δ* = *δ′* = 2*δ″*. Other parameters as in figure 2D of main text : *δ* = 0, 0.1, 0.2, 0.3. *c* = *c′* = 0, *n*_*max*_ = 40. Dotted lines: approximation (9) of main text. The approximation does not work as well as when *δ* = *δ′* = *δ″*.

**Figure S5:**
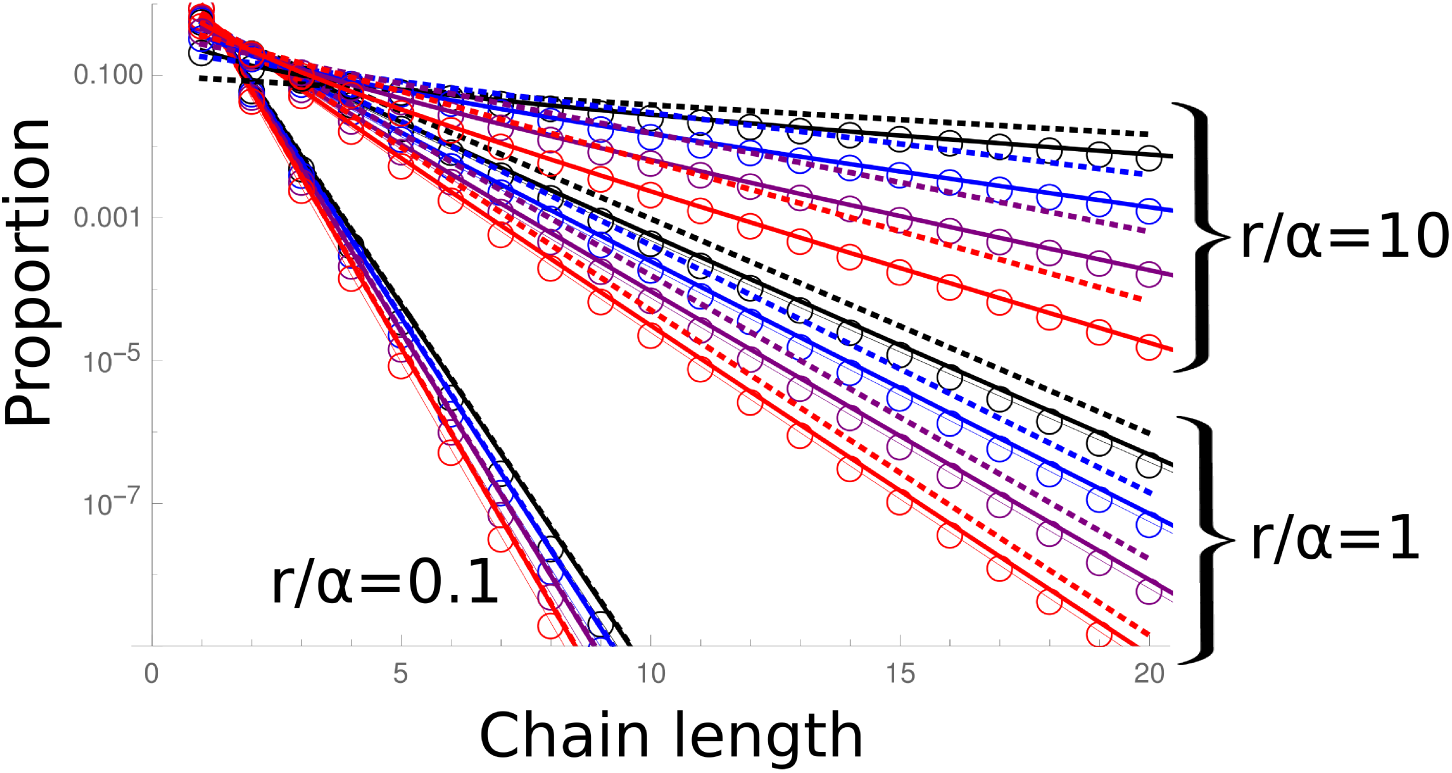
Chain length distribution. Similar to figure 2D of main text. *δ* = *δ′* = *δ″* = 0, 0.1, 0.2, 0.3. Thick solid lines with no markers are for *c* = *c′* = 0, Thin solid lines with *o* are for *c* = 0.2*α, c′* = 0.5*α*. Dotted lines: approximation (9) of main text. There is very little change in the chain length distribution. *n*_*max*_ = 40.

### 4 Chain length distribution with a fixed replication time - approximation

Below, we present in details the assumptions and calculations to obtain the approximation of the chain length distribution when bacteria replicate every *τ*.

We define *n*_*i*_(*t*) the number of chains of length *i* at *t* with *t* taken just before a replication. Assuming *i* even,

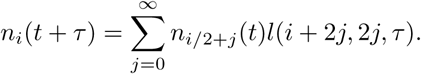

This is because just before a replication, there are *n*_*i/*2+*j*_(*t*) chains of length *i/*2+ *j*. Then, just after the replication, these chains are of length *i* + 2*j*. Time *t* + *τ* is just before the next replication. With probability *l*(*i* + 2*j*, 2*j, τ)*, these chains of length *i* + 2*j* have lost 2*j* bacteria on their edges and are now chains of length *i*. We sum over all the possible *j*. In the long time, *n*_*i*_(*t*) = *Cp*_*i*_ exp(*λt*), with *λ* the long term growth rate, that is such that exp(*λτ)* = 𝒩, with 𝒩the largest eigenvalue of the matrix. Replacing *l*(*i* + 2*j*, 2*j, τ)* by its expression as in equation (11) of the main text, the previous equation leads to:

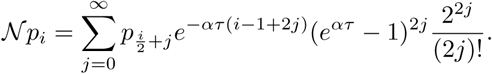

We compare the 1st term of the sum to the rest of the sum. The first term is 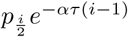, the rest of the sum is:

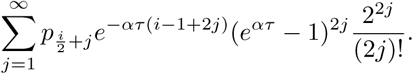

We divide both by *e*^*−ατ*(*i-*^1^)^. Then this is equivalent of comparing *p*_*i/*2_ with:

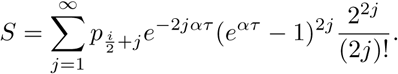

When *ατ* is large, links typically break before the next replication, so there is little cluster formation, and it is thus expected that the chain length distribution decreases fast with *i*, so that for 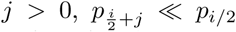. When *ατ* is small, replication is fast compared to the typical time for one link to break. However, for a chain of length *i/*2, *τ* has to be compared to (*i/*2 *−* 1)*/α*, the typical first link breaking time, thus we expect *n*_*i*_ to decrease with *i* for *i* large enough, thus 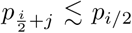 for *j* > 0. We define *B* such as 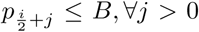 For *ατ* large, *B* ≪ *p*_*i/*2_, and for *ατ* small, if *i* is large enough, *B* ≲*p*_*i/*2_. Then:

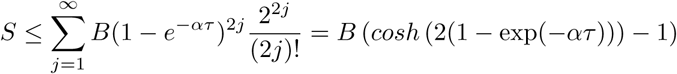

For *ατ* large, (*cosh* (2(1 *−* exp(*-ατ)*)) *−* 1) ≲ *cosh*(2) *−* 1 ≲ 2.7. For *ατ* small, (*cosh* (2(1 *−* exp(*-ατ)*)) *−* 1) ≲ 2(*ατ)*^2^ ≪ 1.

Thus in the case of *ατ* large, *S* is small relative to *p*_*i/*2_ because *S* is smaller than a few units times *B*, with *B* much smaller than *p*_*i/*2_. ln the case of *ατ* small, *S* is small relative to *p*_*i/*2_ because *S* is of the order of (*ατ)*^2^*B*, with *B* of the order of *p*_*i/*2_. Then this justifies the assumption that only the first term of the sum matters:

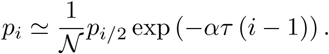

We assume *i* = 2^*k*^, with *k* an integer. This is obviously true only for a very restricted set of *i*, but however this still yields an approximation for how the distribution depends on *i* for large *i*. Then, by recursion,

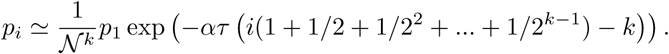

If *i* is large enough, 1+ 1*/*2+ 1*/*2^2^ + *…* + 1*/*2^*k-*1^ ≲ 2. Remembering that *k* was defined as *i* = 2^*k*^, the result is:

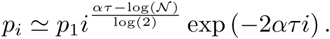

When *ατ* ≫ 1, links typically break before the next replication, thus there is little impact of the clustering on the growth. Consequently, the growth will be close to its value in the absence of clustering, i.e. doubling every *τ*, and thus in this limit 𝒩 = 2:

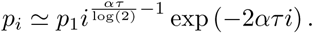

This rough approximation allows to explain the core of the observed distribution.

### 5 Model with force-dependent breaking rate

#### 5.1 Model and equations

A link between bacteria may consist of several sIgA bonds, and the number of bound sIgA may not be exactly the same from one inter-bacteria link to the next, but as sIgA are likely well mixed, many per bacteria and that bacteria are similar to each other, let us assume that link heterogeneity is negligible. The links could break if there was some process degrading the sIgA, but the sIgA are thought to be very stable[6]. Another possible explanation for link breaking is that the antigen get extracted from the bacterial membrane, at a rate which may depend exponentially with the force applied on the link[7] [8]. If the forces are produced by the bacteria themselves(such as by flagella rotation),there are likely to fluctuate on timescales which are short compared to the time between two bacterial replications, and their distribution is likely to be the same for all links, so it would be appropriate to model their effect as a fixed breaking rate, the same for all the links. Another force is the hydrodynamical force exerted by the flow on the bacterial chain.

The flow in the digestive system is complex and not precisely characterized. Longer bacterial chains may also bend and their shape have complex interactions with the flow. Here, we present the simplest model taking into account the forces exerted by the flow on the link breaking rate. We aim to capture the main plausible effects of the flow when the link breaking rate is force-dependent.

**Figure S6:**
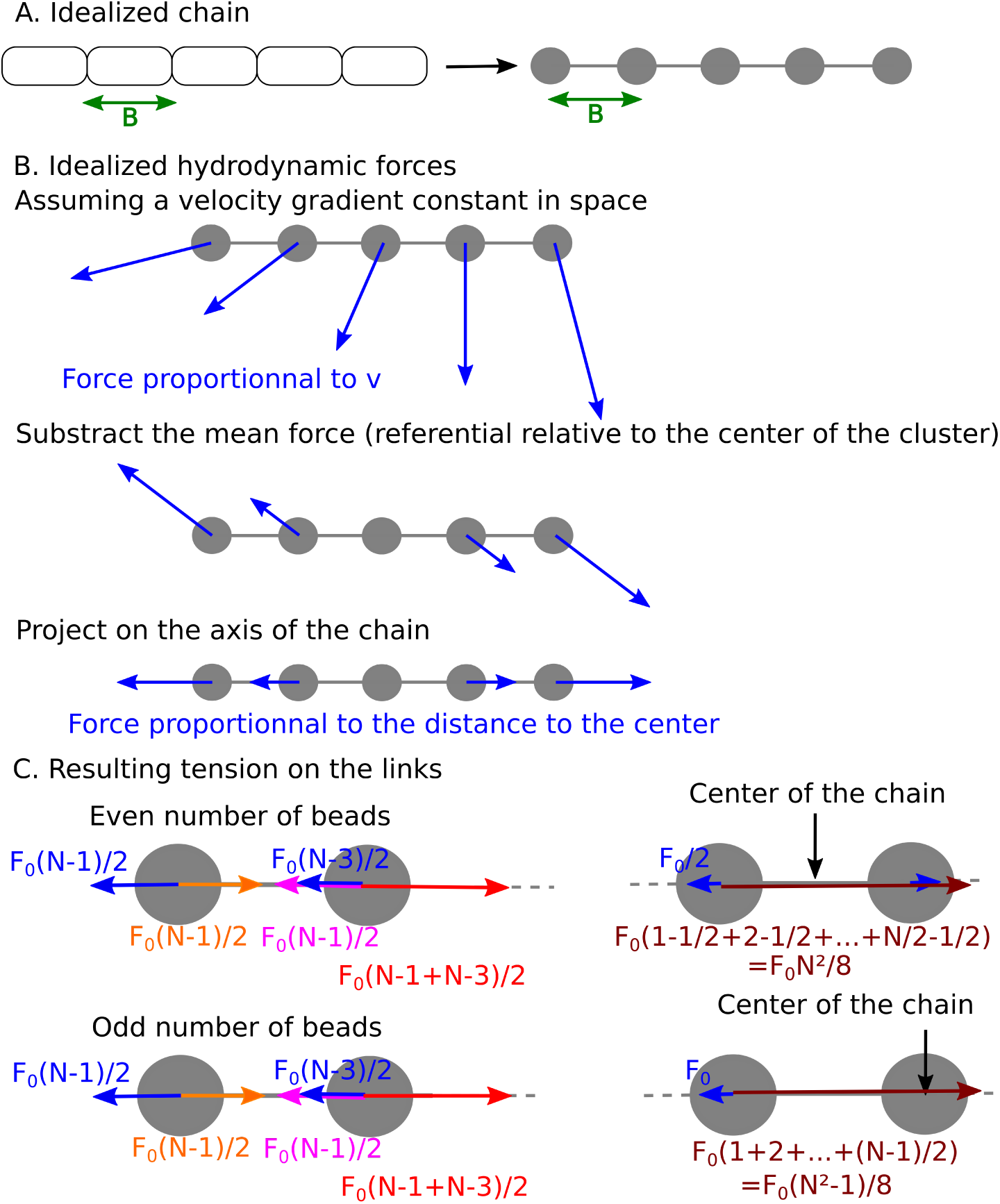
Schematic of the forces applied to the chain. **A** We assume a straight chain of beads with no hydrodynamic interactions between them. B We substract the average force to put ourselves in the referential of the center of the chain, as the total force will translate the whole chain and not impact on the forces on the links. We focus on the forces parallel to the chain that will impact the tension between the links. C Sum of the forces on each bead, for chains with even and odd number of beads.

Let us take a linear chain of *N* bacteria, each of length *B*. Let us approximate it by a rigid chain with beads linked by straight rods of length *B* (pannel A of figure S6). Let us assume that the rods are infinitely thin so they do not interact with the flow, and let us neglect the hydrodynamical interaction between the beads, so they each are subject to the same frictional force for a given fluid velocity, and, given that the typical Reynolds numbers in the digestive tract are relatively low[9], then the viscous force on each bead is proportional to the flow velocity.

Then, let us assume that the velocity gradient in the fluid is constant around the chain. The rationale for this approximation is that the typical scales of the flow are of the order of the centimeter / millimeter (for instance in a mouse, the cecum typical size is in the *cm* range), much larger than typical bacterial chains (the length of one bacteria is about 2*μm*, so even chains of dozens of bacteria remain small compared to the typical flow scale), thus we take a linear approximation of the velocity field in the vicinity of a bacterial chain.

Then, if we take the sum of the forces on the whole chain, it will be equal on *mN* multiplied by the acceleration of the center of mass of the chain, with *m* the mass of each bead. When all the beads move together, there is no force on the links, thus let us take the referential relative to the center of the chain, and subtract the mean force on each bead (panel B of figure S6). Then, there remain forces perpendicular to the axis of the chains, and forces parallel to the axis of the chain. The forces perpendicular to the axis of the chain will make it rotate, and as they are perpendicular, they have no effect on the tension on the rods. Then, let us consider only the forces parallel to the chain.

In the example portrayed here, the chain is elongated. The reverse could happen, but in this case, the chain would likely buckle, and the force applied on the links would be small. The flow varies considerably in time, due to peristaltic motions [10][9]. There would be moments with no force and little breaking, and moments with larger forces and more breaking. The flow due to peristaltic motions changes on time scales short compared to the typical bacterial division time, thus we will assume that periods of low breaking and high breaking rates will be equivalent to an average effective breaking rate. Then let us consider the case of elongation only, as portrayed here.

As we assume here that the velocity gradient is constant, the relative fluid velocity grows linearly with the distance from the center of mass of the chain. Then the force on each bead is equal to *F*_0_ multiplied by the distance to the center divided by *B*. We assume, following [7][8], that the breaking rate is dependent on the force. Thus, we define *α* and *β* such that the breaking rate of a link is *α* exp(*βF/F*_0_) if a force *F* is applied to the link. In the limit of small force, the breaking rate will be *α*, the same for all links, as in the base model. *β* is some constant caracterizing how much the stability of the link is force-dependent.

We can write the force on each bead (pannel C of figure S6). Then, here, because the chain is rigid and straight, the sum of the forces on each bead has to be zero. The tension on the outermost link will simply be equal to the flow force on the outermost bead, i.e. *F*_0_ multiplied by its distance to the center divided by *B*, i.e. (*N* − 1)*/*2 (both for chains of odd and even number of beads). On the next link, the tension has to compensate for the flow force on the second bead, plus the tension applied by the outermost link. Thus the tension on this link is *F*_0_((*N* − 1)*/*2+ (*N* − 1)*/*2 − 1), and so forth (this is analogous to modelling of breaking of polymer chains in elongational flows, as in[11]).

For *N* even, the force on the *j*^*th*^ link starting from the outermost link will be:

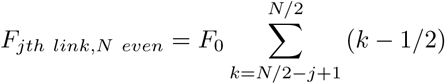

Using *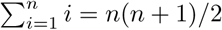*, it can be rewritten as:

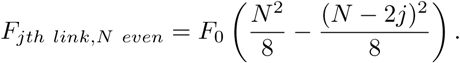

There are two links *j*^*th*^ away from the extremities, for *j* from, to *N/*2 − 1, and one central link, for which *j* = *N/*2. The breaking rate of a given link is *α* exp(*βF/F*_0_) with *F* the total force applied to the link. Then the total breaking rate of one chain of length *N* even is:

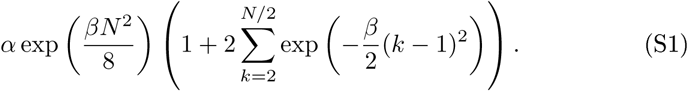

An outermost link of a chain of length *N* + 1 (with *N* even, *N* + 1 is odd) breaks at rate *α* exp(*βN/*2). There are two such links for each chain. This and equation (S1) lead to equation (30) of the main text:

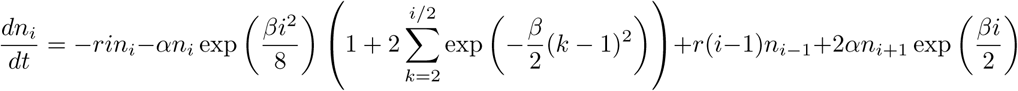

For *N* odd, the force on the *j*^*th*^ link starting from the outermost link will be:

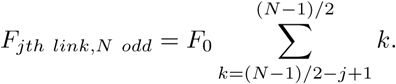

Simiarly to the *N* even case, we can rewrite:

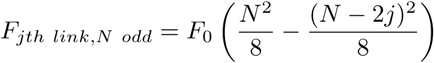

Because of the two sides, there are two links *j* for each chain, for *j* from, to (*N* − 1)*/*2. The breaking rate of a given link is *α* exp(*βF/F*_0_) with *F* the total force applied to the link. Then the total breaking rate of one chain of length *N* odd is:

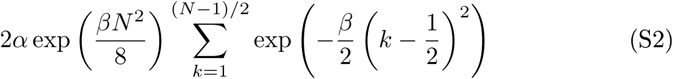

An outermost link of a chain of length *N* +1 (with *N* odd, *N* + 1 is even) breaks at rate *α* exp(*βN/*2). There are two such links for each chain. Then, this and equation (S2) lead to equation (31) of the main text for the evolution in time of the mean number of chains of odd length *i*:

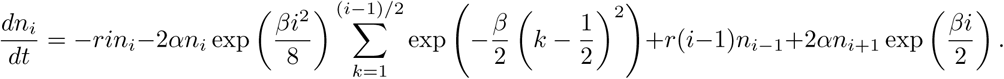

#### 5.2 Additional figure for the force-dependent model: replication rate maximizing the growth rate as a function of *β*

Figure S7 shows that the rate of replication maximizing the growth rate of free bacteria increases exponentially with *β*, which represent the strength of the dependence of the breaking rate on the force applied to the link.

**Figure S7:**
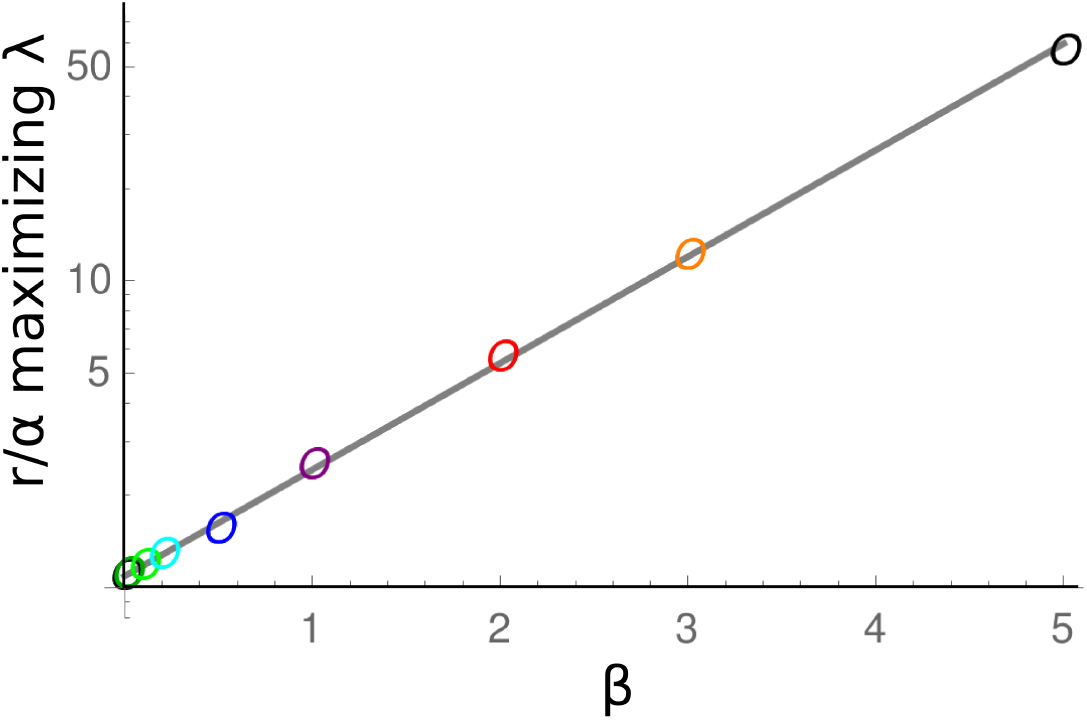
Log of the value of *r/α* maximizing the growth rate in the force-dependent breaking rate model as a function of *β*. The points are numerical maximums, the line is 1.09 × exp(0.8*β*). 1.09 is the value of (*r/α*) maximizing the growth rate for the base model (i.e. for *β* → 0).

#### 5.3 Force-dependent model: approximation for the chain length distribution

We start from equations (30) and (31), and assume that for *t* long enough, *n*_*i*_ ≃ *Cp*_*i*_ exp(*λt*) (with *λ* the largest eigenvalue). Then,

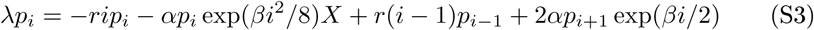

with 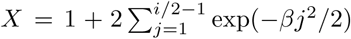 (*i* even) or 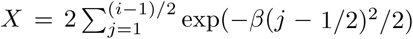 (*i* odd). Let us now determine which terms dominate in this expression.

For *i* large enough, *λ* ≪ *ri*. Thus *λp*_*i*_ is negligible relative to *rip*_*i*_.

For both *i* even and odd, *X* is a converging sum which tends to a finite number when *i* increases. Let us denote its limit 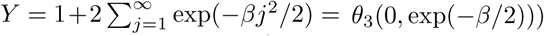 in the even case, and 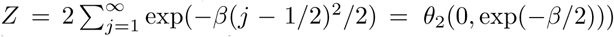 in the odd case, with *θ*_*i*_ the Jacobi Theta functions. Thus, because *β* is positive, for *i* large enough, *ri* ≪ *α* exp(*βi*^2^*/*8)*X*.

The remaining main terms in equation (S3) are:

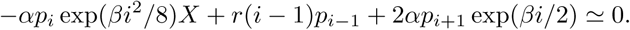

The first term is negative, the two others are positive. Then we have to determine which of *r*(*i* − 1)*p*_*i−*1_ and 2*αp*_*i*+1_ exp(*βi/*2) dominates. If 2*αp*_*i*+1_ exp(*βi/*2) dominates, *αp*_*i*_ exp(*βi*^2^*/*8)*X* ≃ 2*αp*_*i*+1_ exp(*βi/*2), thus *p*_*i*+1_*/p*_*i*_ ≃ exp(*βi*(*i/*8 − 1*/*2))*X*, which for *i* large enough means that the long the chain, the more of it, which would diverge and does not make sense in this system. Thus *αp*_*i*_ exp(*βi*^2^*/*8)*X* ≃ *r*(*i* − 1)*p*_*i−*1_,

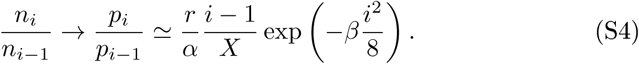

This approximation is valid for large chain sizes. We assume that it is valid for any chain length. As this expression is small and decreasing quickly with increasing *i, p*_1_ will be close to 1. Then, as:

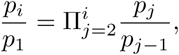

and using the known expression for the sum of the squares 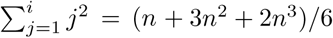 and expression (S4):

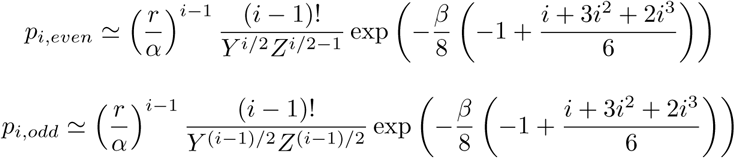

These two equations can be combined, and ultimately lead to:

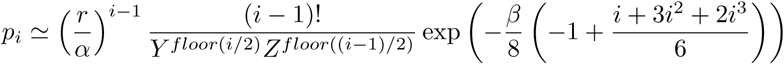

#### 5.4 Additional figure for the force-dependent model: chain length distribution for other values of *r/α*

In panel 2J of the main text, we represented the distribution of chain lengths in the model with force-dependent link breaking rate for *r/α* = 1. In figure S8 we represent the distributions for different values of *r/α*. Overall, the shapes are similar, and the smaller *r/α* is (as well as the larger *β* is), the better the analytical approximation works.

**Figure S8:**
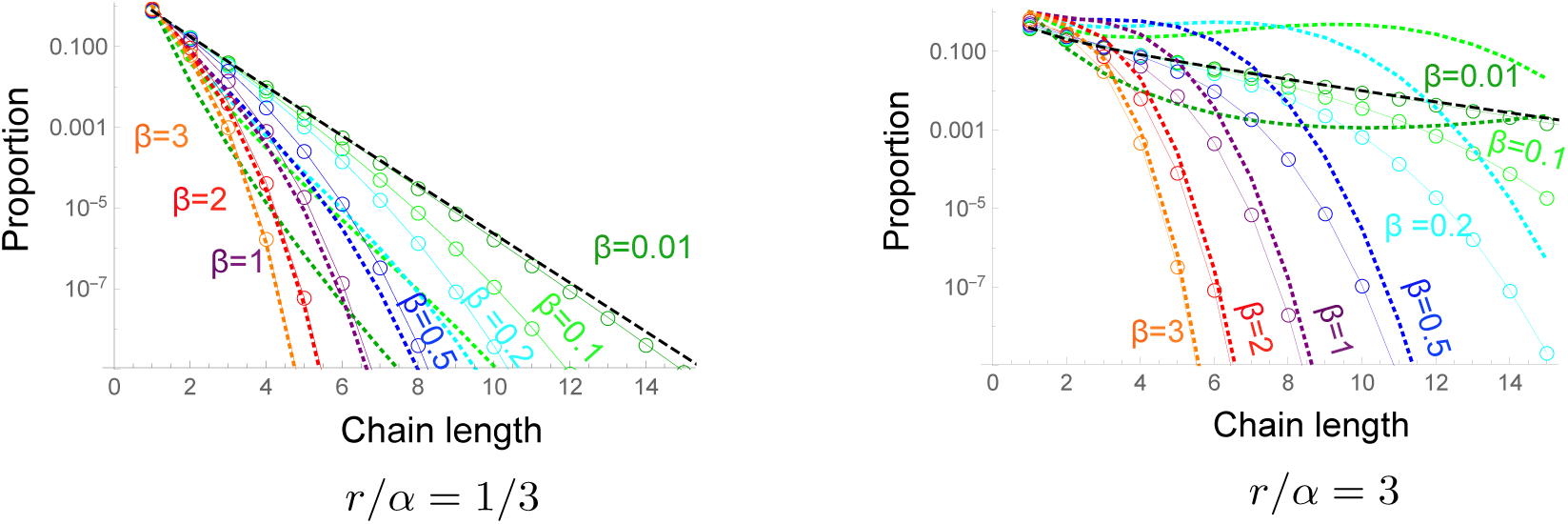
Chain length distribution, as in figure 2J, except for the value of *r/α*: model with force-dependent breaking rates. Each color represents a different *β*: *β* = 0.01 (*n*_*max*_ = 20), *β* = 0.1 (*n*_*max*_ = 15), *β* = 0.2 (*n*_*max*_ = 15), *β* = 0.5 (*n*_*max*_ = 15), *β* = 1 (*n*_*max*_ = 15), *β* = 2 (*n*_*max*_ = 10), *β* = 3 (*n*_*max*_ = 10). The black dashed lines are the numerical results for the base model, equivalent to *β* = 0. The curves for *β* = 0.01 (dark green) are almost overlaid with the curves for *β* = 0. The colored dotted lines the analytical approximation (equation (18) of the main text), and the black dotted line the approximation for the base model (equation (5) of the main text).

### 6 Zoomed-in distribution for all models

Figure S9 represents the zoomed-in chain length distributions of the right panels of figure 2 of the main text.

**Figure S9:**
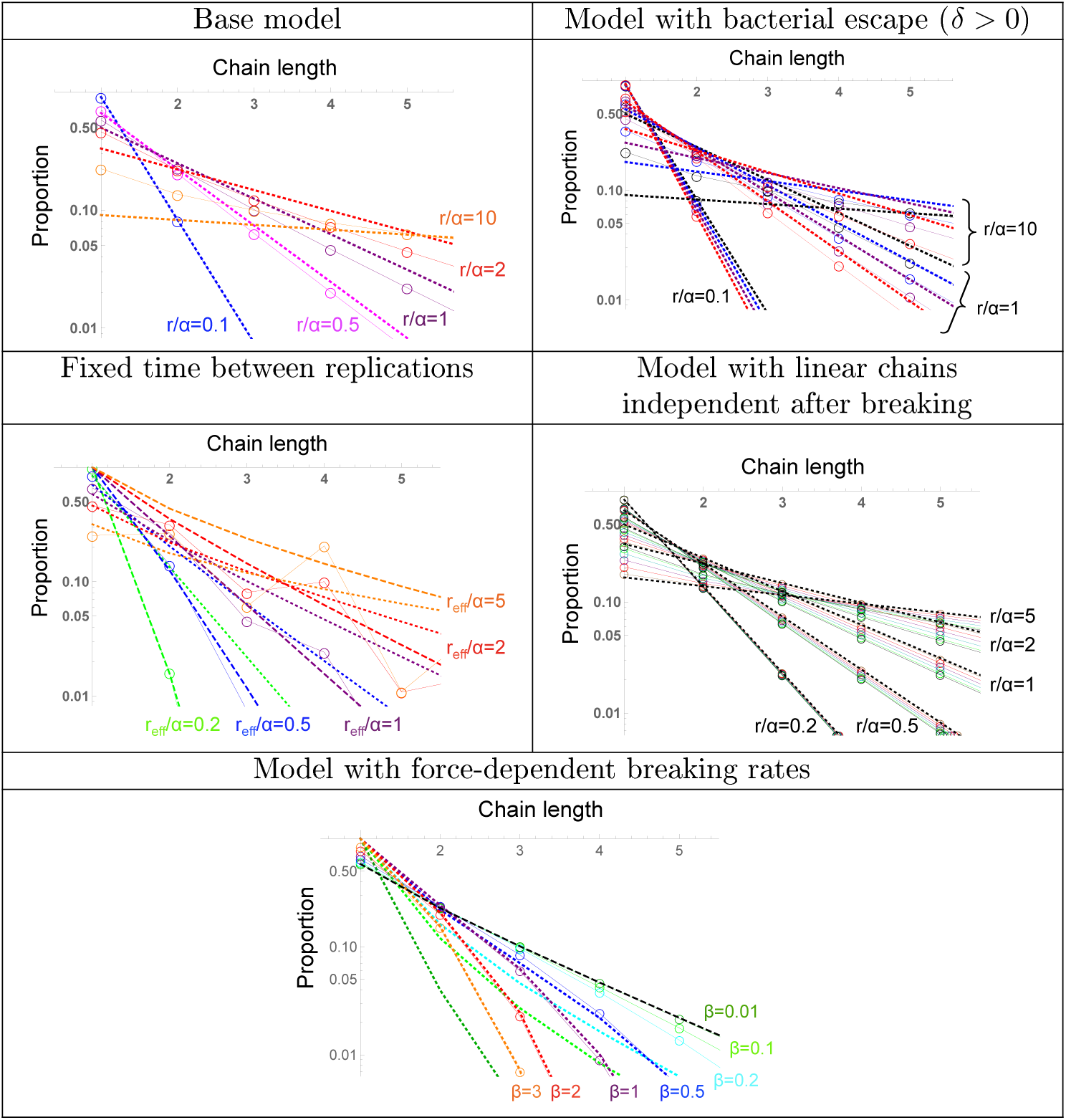
Zoomed-in chain length distributions: everything as in the right panels of figure 2 of the main text, but centered on the head of the distributions. Solid lines and open circles: numerical results. **Base model**: *n*_*max*_ = 40, dotted lines: approximation (5) of the main text (almost overlaid with the numerical results for *r/α* = 0.1). **Model with bacterial escape**: *δ* = *δ′* = *δ″* = **0**,0.1, 0.2, 0.3, *c* = *c′* = 0, *n*_*max*_ = 40. Dotted lines: approximation (9) of the main text. **Fixed time between replications**: *r*_*eff*_ = log(2)*/τ, n*_*max*_ = 32. Approximation (18) (dashed lines), numerical result in the base model (dotted lines). *r/α* = 0.2, 0.5, 1, 2, 5. Model with linear chains independent after breaking: *n*_*max*_ = 100. The dotted black lines are the approximate distribution (27) of the main text for each *r/α*, which is the exact distribution for *q* = 1. The colors represent *q* =*β* = 0, 0.1, 0.2, 0.3, 0.4, 0.5, 0.6, 0.7, 0.8, 0.9, **1**. All curves are almost overlaid for small *r*. **Model with force-dependent breaking rates**: chain length distribution for *r/α* = 1. Each color represents a different *β*: *β* = 0.01 (*n*_*max*_ = 20), *β* = 0.1 (*n*_*max*_ = 15), *β* = 0.2 (*n*_*max*_ = 15), *β* = 0.5 (*n*_*max*_ = 15), *β* = 1 (*n*_*max*_ = 15), *β* = 2 (*n*_*max*_ = 10), *β* = 3 (*n*_*max*_ = 10). The colored dotted lines the analytical approximation (18) of the main text, and the black dotted line the approximation (5) of the main text for the base model.

### 7 Experimental data

#### 7.1 Methods

We perform a new analysis on images that were produced for [4]. We briefly describe below the experiments from which the images were produced, and describe our analysis.

Mice, which were previously vaccinated with a peracetic-acid inactivated S.Typhimurium strain (PA-S.Tm), were pretreated with 0.8g/kg ampicillin sodium salt in sterile PBS. 24h later, mice received 10^5^ CFU of a 1:1 mix of mCherry- (pFPV25.1) and GFP-(pM965) expressing attenuated S. Tm M2702. For imaging, cecum content was diluted gently 1:10 w/v in sterile PBS containing 6 *μ*g/ml chloramphenicol to prevent growth during imaging. 200*μ*l of the suspension were transferred to an 8-well Nunc Lab-Tek Chambered Coverglass (Thermo Scientific) and imaged at 100x using the Zeiss Axiovert 200m microscope. To determine the distribution of bacteria in aggregates, n=25 high power fields per mouse were randomly selected and imaged for mCherry and GFP fluorescence. For some mice, sequential sampling was done, these mice were terminally anaes-thetised and artificially respirated cecum content was sampled by tying off part of the cecum each hour for 3h. More details about the experimental procedures can be found in [4].

We analyzed all the images for the early data points (4 and 5 hours) of experiments starting from a low inoculum (10^5^), to minimize the clustering from random encounters. Only the linear chains were counted. Images are for the red and green fluorescence, so complex clusters with two colors were not counted. The data were analyzed manually. The images are available as supplementary materials:

- images4h.zip contains the images for 3 of the mice only sampled at 4h.
- images4h_others.zip contains the images for the other 4 mice only sampled at 4h.
- images5h.zip contains the images of the mice only sampled at 5h
- imagesseq4h.zip contains the images at 4h of mice sampled sequentially
- imagesseq5h.zip contains the images at 5h of mice sampled sequentially

#### 7.2 Results

For linear chains, we obtained the length distribution detailed in table 1 and shown on figure 4 of the main text.

**Table 1:**
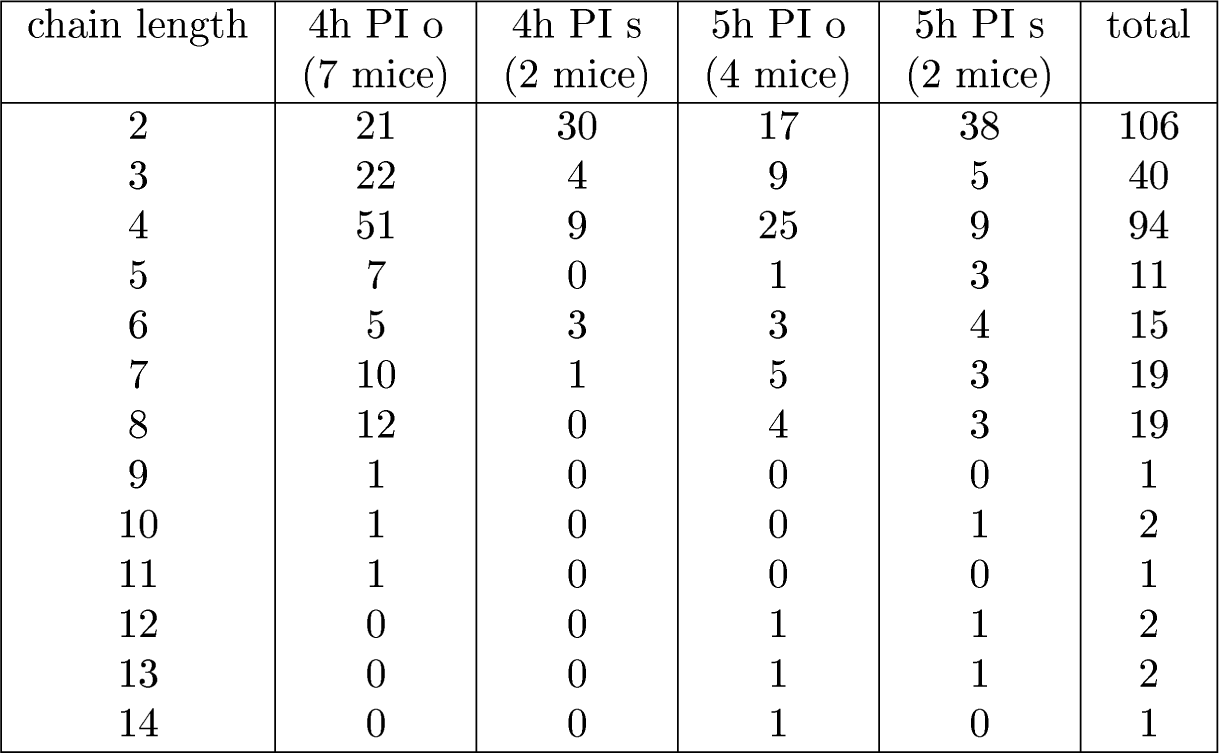
Table of the linear chains counted on the images from several experiments, either with mice sampled once (o), or with mice sampled sequentially (s).

| chain length | 4h PI o<br>(7 mice) | 4h PI s<br>(2 mice) | 5h PI o<br>(4 mice) | 5h PI s<br>(2 mice) | total |
| --- | --- | --- | --- | --- | --- |
| 2 | 21 | 30 | 17 | 38 | 106 |
| 3 | 22 | 4 | 9 | 5 | 40 |
| 4 | 51 | 9 | 25 | 9 | 94 |
| 5 | 7 | 0 | 1 | 3 | 11 |
| 6 | 5 | 3 | 3 | 4 | 15 |
| 7 | 10 | 1 | 5 | 3 | 19 |
| 8 | 12 | 0 | 4 | 3 | 19 |
| 9 | 1 | 0 | 0 | 0 | 1 |
| 10 | 1 | 0 | 0 | 1 | 2 |
| 11 | 1 | 0 | 0 | 0 | 1 |
| 12 | 0 | 0 | 1 | 1 | 2 |
| 13 | 0 | 0 | 1 | 1 | 2 |
| 14 | 0 | 0 | 1 | 0 | 1 |

Given the bumpy shape of the experimental distribution, we chose to fit the data with the fixed replication time model for the figure 4 of the main text. In this model, the only adjustable parameter is *r*_*eff*_ */α*. For a given *r*_*eff*_ */α*, we obtain the theoretical chain length distribution *p*_*i*_ by numerical resolution of the equations. As the data concerns only chains of length 2 and longer, we renormalize this distribution as *p*_*i*_*/*(1 *- p*_1_). Then, given that we observe a total of *N*_*exp*_ = 313 chains, for the theoretical process the probability to observe *k* chains of length *i* is *Poisson*(*k, λ* = *N*_*exp*_*p*_*i*_*/*(1 *- p*_1_)) = *λ*^*k*^ exp(*-λ*)*/k*!. Naming *k*_*i*_ the number of observed chains of length *i*, we then take the value of *r*_*eff*_ */α* which maximizes:

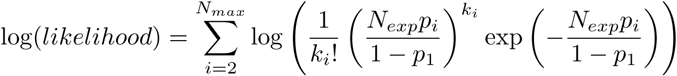

Given that we compute numerically the theoretical distribution, we have to use a finite *N*_*max*_, whereas in theory it should be taken as infinite. In practice, the theoretical values for the proportions of long chains become quickly very small, and thus the probability to observe chains of such lengths is very small for any *r*_*eff*_ */α*. Thus in order to fit *r*_*eff*_ */α, N*_*max*_ does not matter, provided that it is taken large enough. We have tried *N*_*max*_ = 16 and 32, and we obtain the same fitted value *r*_*eff*_ */α* = 4.1. If we were to look for a value with more significant digits, this choice for *N*_*max*_ would matter more.

To quantify our impression that there are fewer long chains observed than expected, we performed the following calculations. Taking *r*_*eff*_ */α* = 4.1 and *N*_*exp*_ = 313, the expected number of chains of length 15 or longer is 3.7, whereas none is observed, which with a Poisson assumption has a probability 0.025 to occur. This probability seems low. Either this is a low probability but still happened (and if we look at a bit shorter chains, the expected average number of chains of length 9 and longer is 11.7, and 9 of them are actually observed, which is relatively close); or there is some process limiting the number of long chains. There are two main possibilities for the number of long chains to be limited: there could be an experimental bias limiting the observation of long chains (see discussion below); or there could be some force-dependence of the breaking rates, which would effectively act as a cut-off for the chain length (see figure 3 of main text), as in this case, breaking rates increase considerably with chain length.

#### 7.3 Discussion

The data may be biased. The mass of one bacterium is about one pg, and its density is about 10% more than the water density[12, 13], the thermal energy at ambient temperature is of the order of 4.10^*−*21^*J*, and gravity *g* is of the order of 10*m/s*^2^, thus thermal fluctuations will lift an individual bacterium by typically 4 *μ*m higher than the bottom. Thermal fluctuations will have two effects:

- The average height of the center of gravity of chains will decrease with their length. This is confocal microscopy, which typical optical section is less than 1*μ*m, focused close to the cover slip. This may bias the distribution by missing smaller chains.
- Longer chains are not rod-like, their shape fluctuate. It is apparent on the microscopy images that parts of long chains may get out of focus. The longer the chain, the less likely that it is entirely in the focus, and thus chains will look smaller than they are.

We focus on the chain length distribution because this quantity is more easily accessible by experimental measurements, at the end of an experiment. Comparing models and experiments enables to check whether the data is compatible with a process of growing and breaking of clusters; and determine which specific model is closest to the data. However, some models cannot be distinguished, no matter how much data is available for the chain length distribution. For example in the model with bacterial escape and the model where chains can remain independent after breaking, there are two parameters to fit (*r/α* and *δ*, or *r/α* and *q*). It is likely that fitting would mainly select a value for *r/α*, since the distribution does not depend much on the second parameter in both cases. These models could not be distinguished from the base model. On the other hand, models with different distribution shapes - either in the force-dependent model or in the fixed division time one-could be distinguished, provided that the bias can be overcome, and that more data can be collected.

We could fit the fixed replication time model to the data, and this strengthened our hypothesis that the chains are generated by a process of enchained growth and link breaking. However, there is somewhat less long chains observed than expected (especially in the range of lengths 14 to 16). One possibility could be that the breaking rate is force dependent. If we had 10 times more of unbiased data, we could answer whether there really is a deficit of longer chains. If there is indeed a deficit of longer chains, then we should combine the model of force-dependent breaking rates with the model with fixed replication time, to be able to make quantitative comparisons. To be more effective, comparison would likely require more data, as there would be two free parameters, *r/α* and *β*. If there is no deficit of longer chains in the range up to 16, then the simple model with fixed replication time predicts that we would get access to the distribution up to length 24 (and be at the limit for lengths 28 and 32) with 100 times more data than in current experiments (see figure S10). Thus overall we would need at least 10 times and likely 100 times more data for a more quantitative assessment.

**Figure S10:**
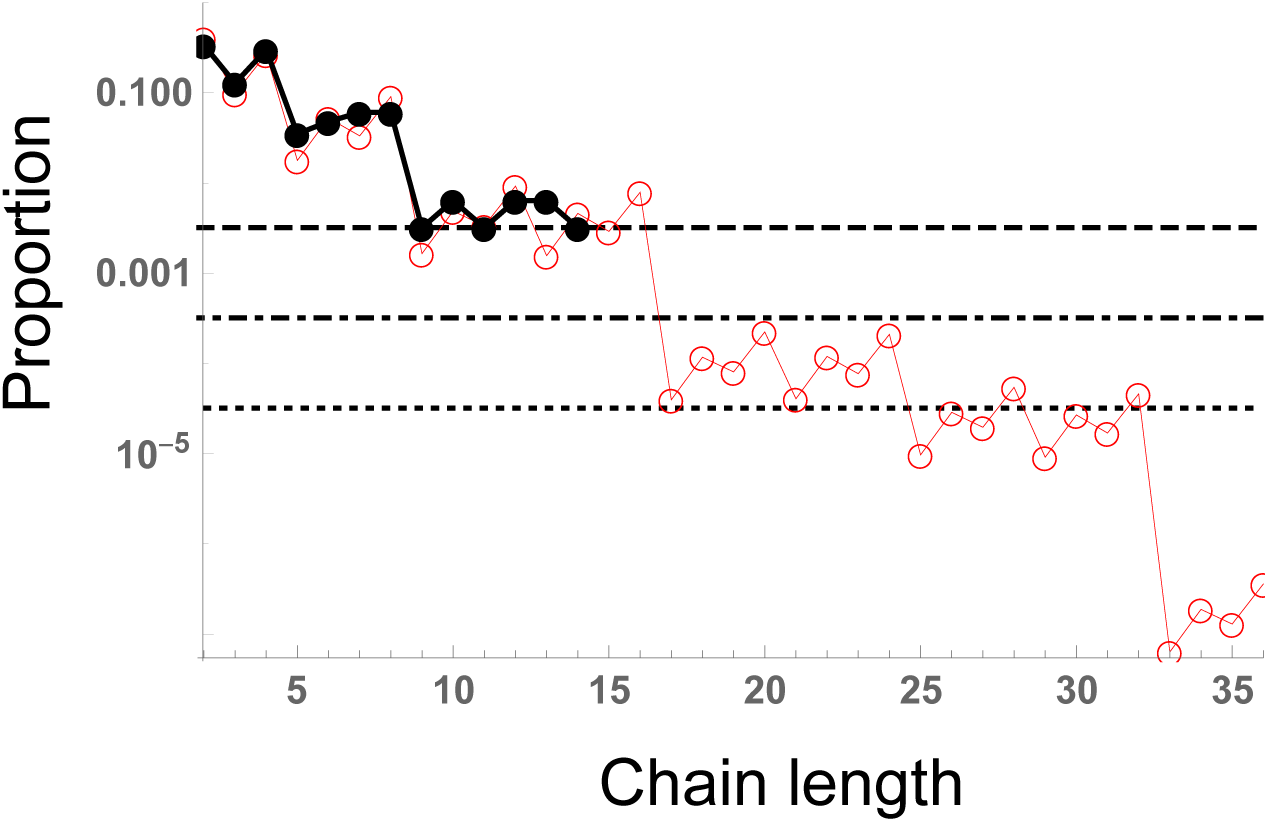
Comparison of chain length distribution for the experimental data (black) and the model with fixed replication time (red). This figure is as figure 4 of main text, but extended up to length 36. The black dashed line represents limit of one chain in the data. The black dot-dashed line represents what this limit would be if there was 10 times more data, and the black dotted line represents what this limit would be if there was 100 times more data. *r*_*eff*_ */α* = 4.1 (with *r*_*eff*_ */α* = log(2)*/*(*ατ)*).

Increasing the amount of data would not necessarily require to sacrifice more mice, but merely to take more images for each cecum content. The challenge would be to do so with no bias, and with very standardized conditions so that the images are taken in conditions close enough so as to automate the chain detection and length count.

It would be also very useful if there would be ways to estimate the breaking rate in independent experiments, for instance injecting (without breaking them nor perturbing the system) chains of non-replicating bacteria of controlled length, and measuring how the length distribution changes over time. Then, as the replication rate can be estimated by other measures (dilution of non-replicating plasmids), we could get an estimate of the replication rate over the breaking rate, which would considerably constrain the fitting of the chain length distribution, and thus give more strength to the conclusions achieved.

